# Persistent Immune Dysregulation during Post-Acute Sequelae of COVID-19 is Manifested in Antibodies Targeting Envelope and Nucleocapsid Proteins

**DOI:** 10.1101/2025.08.18.670908

**Authors:** Marcin Kwissa, Manikannan Mathayan, Satyajeet S. Salunkhe, Velavan Bakthavachalam, Zijing Ye, Mark A. Sanborn, Samantha Condo, Aditi Upadhye, Athulith Nemakal, Justin M. Richner, Sanjib Basu, Richard M. Novak, Jeffrey R. Jacobson, Balaji B Ganesh, Martha Cerda, Paul J. Utz, Jerry A. Krishnan, Bellur S. Prabhakar, Jalees Rehman

**Author notes:** Please address correspondence to: B.S.P or J.R. These authors contributed equally (Marcin Kwissa, Manikannan Mathayan).

## Abstract

Post-Acute Sequelae of SARS-CoV-2 infection (PASC) syndrome or “Long COVID” represents a widespread health challenge that necessitates the development of novel diagnostic approaches and targeted therapies that can be readily deployed. Immune dysregulation has been reported as one of the hallmarks of PASC, but the extent of PASC immune dysregulation in patients over time remains unclear. We therefore assessed SARS-CoV-2-specific antibody responses, peripheral immune cell profiles, autoantibody profiles and circulating cytokines for up to 6 months in participants with a SARS-CoV-2 infection who either convalesced or developed PASC. Compared to convalescent, PASC participants with a broad range of PASC phenotypes exhibited persistently elevated IgG titers for SARS-CoV-2 Envelope and Nucleocapsid proteins over the 6 months of study duration. In contrast, the IgG responses to Spike protein were significantly lower in the PASC cohort with predominantly IgG1 and IgG3 class-switched bias. Using CyTOF analysis, we show elevated numbers of circulating T follicular helper cells (cTFH) and mucosa-associated invariant T cells (MAIT), which also correlated with high anti-Envelope IgG titers. Persistent immune activation was accompanied by augmented serum cytokine profiles with LIF, IL-11, Eotaxin-3, and HMGB-1 in PASC participants, who also demonstrated significantly higher rates of autoantibodies. These findings highlight the persistence of immune dysregulation in PASC, underscoring the need to explore targeted therapies addressing viral persistence, dysregulated antibody production, and autoimmunity.

## Introduction

Post-Acute Sequelae of SARS-CoV-2 infection (PASC), or “Long COVID”, is a debilitating chronic disease for which there are currently no targeted therapies available and which highlights the enduring impact of acute SARS-CoV-2 infection on human health.^1^. PASC is a syndrome comprised of multiple endotypes or lingering disease manifestations such as excessive fatigue, “brain fog”, dysautonomia, and additional neurological, cardiovascular, musculoskeletal, or respiratory complications ^2–5^. The clinical manifestations vary in duration and severity and affect multiple organ systems, with considerable inter-individual variability^6,7^. Due to the heterogeneity of PASC manifestations and the breadth of the symptoms, diagnosing PASC and distinguishing it from other chronic diseases with congruent symptoms has emerged as a major challenge in clinical practice.

Persistent immune dysregulation is emerging as a fundamental pathogenic mechanism of PASC, and characterizing such immune dysregulation holds great promise because it allows for the development of specific diagnostic biomarkers as well as targeted therapeutics. Several studies have reported sustained alterations in the innate immune system, including prolonged activation of the complement cascade^8,9^ and continued stimulation of innate immune cells of myeloid origin such as monocytes, macrophages^10–12^ and neutrophils^13–15^ as well as natural killer (NK) cells^16,17^. Additionally, dysregulation within the adaptive immune compartment has also been observed, characterized by exhausted T cell phenotypes and aberrant B cell activation^12,16,18,19^. Together, these immune abnormalities suggest that a failure to fully resolve the inflammatory response to the initial SARS-CoV-2 infection may underlie the persistent symptoms observed in individuals with PASC. Additionally, autoimmune responses have also been associated with PASC pathology, such as the presence of autoantibodies (Abs) targeting a wide range of self-antigens, including components of key immunological pathways. Some studies have identified elevated levels of auto-Abs targeting interferons^20,21^, chemokines^22^ or nuclear antigens^23^ in subsets of PASC patients. The observed heterogeneity suggests that such auto-Abs, when present, may reflect underlying predisposition, prior immune priming, or secondary consequences of sustained inflammation, rather than a unifying etiological feature of PASC.

More recent studies suggest that the persistence of SARS-CoV-2 in various tissues may also contribute to the pathogenesis of PASC because lingering SARS-CoV-2 RNA and proteins in tissues could potentially sustain immune activation. Multiple studies have detected viral RNA and proteins in tissues such as the lungs, gastrointestinal tract, and brain months after the acute phase of infection^24–28^. Additionally, disruption of mucosal barrier integrity, particularly in the gut and lungs, has been reported in PASC, raising the possibility of viral persistence promoting microbial translocation that could further perpetuate systemic inflammation and immune imbalance ^4,14,29^.

In addition to the complexity of the immune dysregulation and heterogeneity of PASC, the severity of the symptoms can wax and wane over time, thus creating a challenge for developing diagnostic biomarker panels and choosing optimal temporal windows for therapeutic interventions. Longitudinal studies with serial sampling are therefore required to determine whether immune dysregulation is transient, progressive, or persistent. To date, comprehensive multi-timepoint studies in well-characterized cohorts remain limited, leaving critical questions about temporal immune dynamics in PASC unanswered. Documenting viral persistence in tissues would be valuable for diagnostics and for choosing which patients would benefit from antiviral therapies, but obtaining tissue samples from patients often requires invasive studies that are not feasible for longitudinal analysis. As a result, conclusions regarding viral persistence remain difficult to study non-invasively, and several studies have failed to detect SARS-CoV-2 RNA or protein in blood, stool^30^, or tissue samples from individuals with PASC ^18,31,32^. As an alternative approach, the adaptive immune system may serve as an indirect biosensor of persistent viral antigens and could be assessed by non-invasive means. Long-lived memory B and T cells can retain antigen specificity, and the detection of sustained antibody responses to viral antigens such as nucleocapsid or envelope proteins that are not used in COVID-19 vaccines may offer indirect evidence of ongoing viral antigen exposure and potential viral persistence.

We therefore studied immune dysregulation and antibody responses over time in participants with PASC and those who recovered fully from COVID-19. We show a marked persistence in SARS-CoV-2–specific antibody responses, particularly elevated IgG to envelope and nucleocapsid proteins, along with altered immune cell profiles including increased MAIT and cTFH cells in PASC participants. PASC was also associated with a proinflammatory cytokine environment and the emergence of autoantibodies, suggesting persistent immune activation and dysregulation.

## Results

### Elevated IgG response against SARS-CoV-2 Envelope and Nucleocapsid in PASC

We studied thirty participants who enrolled in the parent NIH RECOVER Adult Cohort Study during the post-acute phase of a SARS-CoV-2 infection. Details about the design of the Adult Cohort Study are available elsewhere^33^. This report examines peripheral blood samples from 30 participants (20 with PASC, and 10 participants who were convalescent (CONV) without any persistent PASC symptoms). Participants in the PASC group presented with different symptoms, including cardio-pulmonary (n=3), fatigue/malaise (n=2), neuropsychiatric (n=1), dizziness (n=4), dysautonomia (n=4) or indeterminate PASC symptoms (n=6), which showcased the variety of global PASC manifestations (**Table 1**), mirroring the heterogeneity seen in a typical PASC patient population ^7,34,35^. To study possible longitudinal shifts in immune dysregulation, peripheral blood samples were obtained at baseline during enrollment into the parent study (Timepoint 0 or T0, on average 42 days post infection), approximately 2-3 months later (T1) and the final timepoint after an additional 2-3 months (T2). We conducted a comprehensive analysis of peripheral immunological signatures in the blood by testing serum antibody repertoire against multiple SARS-CoV-2 antigens, immunophenotyping peripheral blood mononuclear cells (PBMC) using multiparametric CyTOF analysis, analyzing circulating cytokines as well as autoantibodies and the plasma across all time points **(Figure 1A**).

**Figure 1.**
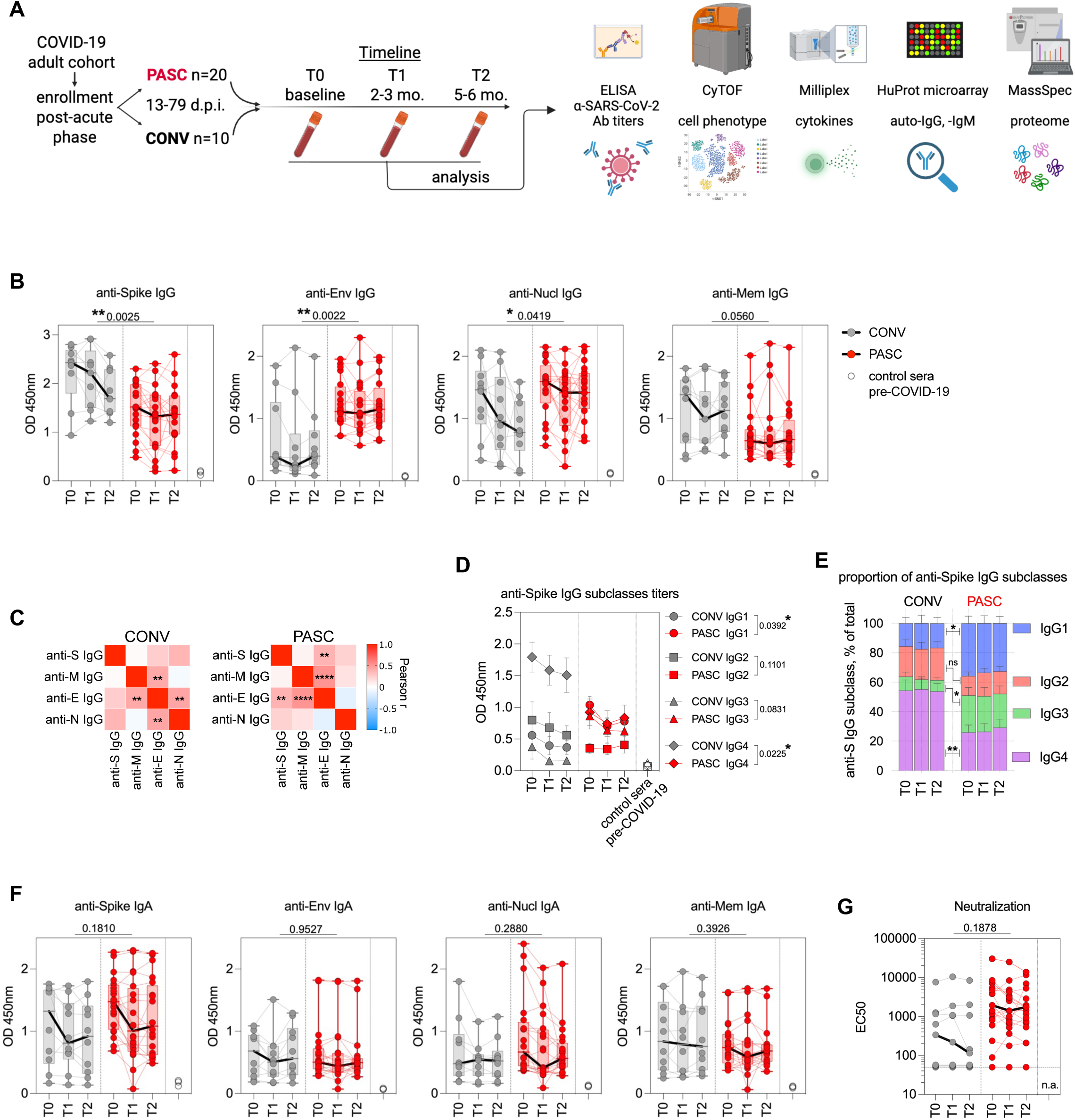
Elevated anti-Envelope and anti-Nucleocapsid IgG titers in PASC participants. (**A**) Experimental design. (**B**) Total IgG titers specific for Spike, Envelope (Env), Membrane (Mem) and Nucleocapsid (Nucl) as measured by ELISA at T0, T1 and T2 collection timepoint. (**C**) Heat maps with correlation matrix of IgG titers (Log10) for distinct SARS-CoV-2 proteins Spike (S), Envelope (E), Membrane (M) and Nucleocapsid (N). Pearson r-values are calculated for titers at T0, T1 and T2 timepoints for CONV (left) and PASC (right) cohorts. (**D**) Titers of IgG1, IgG2, IgG3 and IgG4 specific for Spike. Symbols represent mean titer values at each timepoint (+/-SEM) in CONV (grey) and PASC (red) with distinct symbols for each IgG subclass. Numbers in figure legend are p values and indicate difference in titers between CONV and PASC. (**E**) Proportions of each of the anti-Spike IgG subclass represented as mean % of total IgG response at each timepoint. (**F**) Total IgA titers specific for Spike, Env, Mem and Nucl as measured by ELISA at T0, T1 and T2. (**G**) SARS-CoV-2 neutralization titers are represented as EC50. In **B**, **D** and **F** the Y-axis represents ELISA OD values, graphs show Tukey analysis with min-max, each dot represents individuals in PASC (red), CONV (grey) cohorts connected by color-coded lines or Pre-COVID-19 control sera (empty circles), bar-connecting black lines show medians at distinct timepoints. In **B, D, F** and **G** numbers on the top of each graph show p values between experimental groups calculated for all timepoints by 2-way ANOVA. Black (*) indicate significant difference *p < 0.05, **p < 0.01, ****p < 0.0001. n.s.-not significant (p > 0.05).

**Table 1.** Demographic information of PASC and CONV patients in the COVID-19 cohort.

|  | **CONV** | **PASC** |
| --- | --- | --- |
| **participants, n** | 10 | 20 |
| **biological sex, Female n (%)** | 5 (50%) | 18 (90%) |
| **age, mean (range)** | 48.5 (24-79) | 45.4 (25-75) |
| **race and ethnicity, n** | White, 8 | White, 15 |
|  | Black or African American, 1 | Black or African American, 2 |
|  | Asian, 1 | Hispanic, Latino, or Spanish, 1 |
|  |  | Multiple, 2 |
| **PASC symptom, n** |  | Dysautonomia, 4 |
|  |  | Fatigue-Malaise, 2 |
|  |  | Dizziness, 4 |
|  |  | Neuropsychiatric, 1 |
|  |  | Cardio-Pulmonary, 3 |
|  |  | undefined, 6 |
| **Vaccinated, n (%)** | 10 (100%) | 20 (100%) |

|  | CONV | PASC |
| --- | --- | --- |
| participants, n | 10 | 20 |
| biological sex, Female n (%) | 5 (50%) | 18 (90%) |
| age, mean (range) | 48.5 (24-79) | 45.4 (25-75) |
| race and ethincity, n | White, 8 | White, 15 |
|  | Black or African American, 1 | Black or African American, 2 |
|  | Asian, 1 | Hispanic, Latino, or Spanish, 1 |
|  |  | Multiple, 2 |
| PASC symptom, n |  | Dysautonomia, 4 |
|  |  | Fatigue-Malaise, 2 |
|  |  | Dizziness, 4 |
|  |  | Neuropsychiatric, 1 |
|  |  | Cardio-Pulmonary, 3 |
|  |  | undefined, 6 |
| Vaccinated, n (%) | 10 (100%) | 20 (100%) |

We quantified serum IgG titers of antibodies targeting key SARS-CoV-2 structural proteins such as the spike, envelope (Env), nucleocapsid (Nucl), and membrane (Mem) by ELISA (**Figure 1B-E**). To validate the antibody specificity, pre-COVID-19 sera (n=3) were included as negative controls in all assays. The total IgG responses to Env protein were significantly (p=0.0022) elevated over time in PASC with 1.98-fold higher mean OD450 value of anti-Env IgG when compared to CONV individuals across all timepoints (**Figure 1B**), regardless of the symptom manifestations (**Figure S1**). The IgG titers against the Nucl protein were also significantly higher (p=0.0419) in PASC individuals. In contrast, the IgG response to Spike protein was significantly reduced (p=0.0025) in individuals with PASC compared to the CONV cohort across all time points (**Figure 1B**). We also observed a trend in the reduction of Mem-specific IgG titers among PASC individuals over time, when compared to the CONV control cohort. Titers of total serum IgG response specific for Env protein correlated significantly with anti-Mem and anti-Spike IgG antibody response in the PASC cohort (**Figure 1C**). The viral Spike protein is the target antigen of the anti-SARS-CoV-2 humoral response commonly used in the majority of COVID-19 vaccines ^36^. Therefore, to better characterize the immunological profile of Spike-specific IgG responses, we evaluated longitudinal levels of IgG1, IgG2, IgG3, and IgG4 subclasses in both PASC and CONV cohorts (**Figure 1D, E**). PASC participants exhibited significantly higher titers of Spike-specific IgG1 compared to CONV (p = 0.0392), indicating a sustained Th1-biased response. We observed a similar trend for IgG3, with elevated levels in PASC over time (**Figure 1D and E**). In contrast, CONV participants demonstrated a higher titer of IgG4 (p = 0.0225), consistently across all time points, suggesting a more tolerogenic or regulatory profile following recovery from COVID-19 (**Figure 1D**) ^37–39^. In fact, in CONV the proportion of IgG4 within all IgG specific to Spike was on average at 54.5% (+/- 7%) among all IgG subclasses (**Figure 1E**), showcasing a class-switch when compared to IgG4 in PASC (IgG4=27.2%).

We also tested the Spike-, Env-, Nucl- and Mem–specific titers of IgA in plasma (**Figure 1F**). We noted a trend toward increased levels of anti-Spike IgA responses in the PASC cohort, however, there was no significant difference between experimental groups. None of the other tested SARS-CoV-2 antigens showed substantial differences between the experimental groups over time (**Figure 1F**). Virus neutralization titers in plasma varied markedly among individuals with titers ranging from below detection levels up to 3x10^4^ of IC50 units (**Figure 1G**). We observed a mild increase in the mean neutralization activity against the SARS-CoV-2 infection in sera from the PASC group, however this trend was not significantly different from the mean neutralization titer in the CONV group. Our data thus indicate a distinct shift in SARS-CoV-2 antigen specificity of the humoral IgG response with augmented titers against Env and Nucl proteins in PASC participants. Additionally, anti-Spike IgG titers indicated differential isotype skewing, favoring the pro-inflammatory IgG1/IgG3 subclasses in PASC, in contrast to the IgG4- biased response in convalescent individuals.

### Mucosal immunoglobulin signature in PASC plasma

Next, we used mass spectrometry analysis to characterize the total immunoglobulin (Ig) content in the plasma proteome from PASC and CONV individuals (**Figure 2**). A heatmap of Z-scores representing the relative abundance of Ig proteins in plasma revealed a consistent increase in total Ig protein levels in individuals with PASC compared to convalescent controls (**Figure 2A**). Across all detected Ig peptides and at all longitudinal timepoints analyzed, the PASC group exhibited higher normalized expression, suggesting sustained or dysregulated antibody production. Next, we compared quantities of the constant regions of heavy chains to evaluate total subsets of class-switched, circulating immunoglobulins and their subclasses. Individuals with PASC showed persistently elevated levels of IGHM but also IGHA1 and IGHA2 compared to CONV (**Figure 2B**). The total protein spectrum counts of each of the IgM, IgA1, and IgA2 were significantly more abundant (p=0.0009, p=0.0026, and p=0.004, respectively) in PASC participants when compared to CONV over time. Additionally, J chain protein fragments, which link monomers of IgM and IgA to form polymers, were also highly expressed in PASC (p=0.0028). In contrast, we did not observe differences in the abundance of IGHG1, IGHG2, IGHG3, or IGHG4 between the two groups at either time point. These findings suggest a selective enrichment of mucosal and early-phase antibody isotypes in PASC, potentially indicative of ongoing or dysregulated immune activity, while class-switched IgG subtypes remain stable.

**Figure 2.**
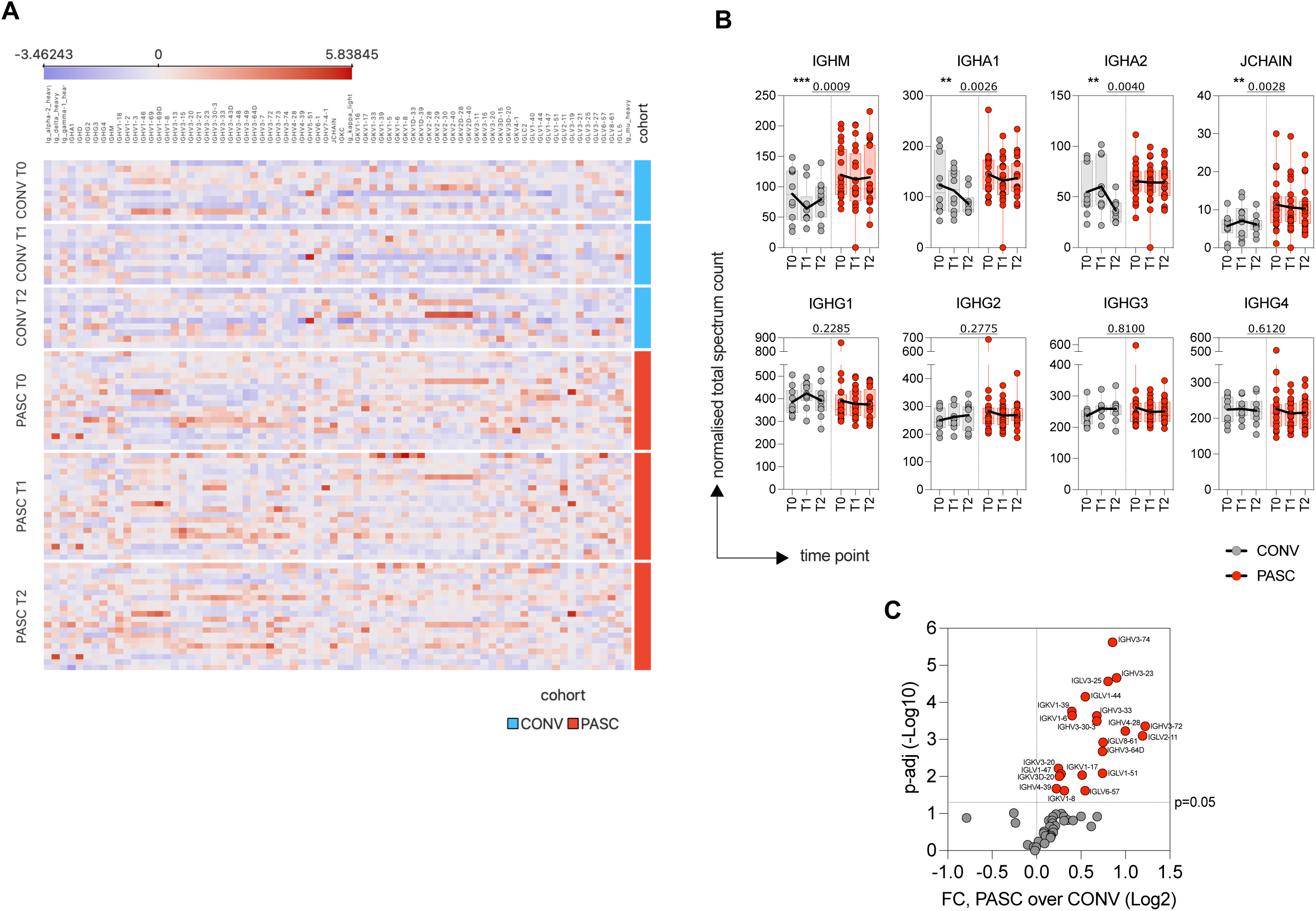
Elevated mucosal immunoglobulin abundance in PASC plasma proteome. Total immunoglobulin content in plasma was analyzed by mass spectrometry. **(A)** The heat map displays Z-scores representing the total abundance of each immunoglobulin isotype or variable region as detected by mass spectrometry-based proteomic analysis. Columns indicate Ig protein annotations; rows represent individual patients from CONV (blue) and PASC (red) cohorts at distinct timepoints as listed on the left. (**B**) Normalized total spectrum count values of heavy chain constant regions: IGHM, IGHA1, IGHA2 and JCHAIN (top row) and IGHG1, IGHG2, IGHG3 and IGHG4 (bottom row) in plasma of PASC and CONV overtime. (**C**) Vulcano plot of intensity values of immunoglobulin variable regions for IGHV, IGL and IGK. Dashed line at y-axis represent an adjusted p-value cutoff of 0.05. Significantly changed cytokines are marked in red. In **B** graphs show Tukey analysis with min-max, each dot represents individuals in PASC (red), CONV (grey) cohorts, bar-connecting black lines show means at distinct timepoints. Numbers on the top of each graph show p values between experimental groups calculated for all timepoints by 2-way ANOVA. Black (*) indicate significant difference **p < 0.01, ***p < 0.001.

We further evaluated potential changes in the Ig heavy, kappa, and lambda variable chains in plasma of PASC and CONV cohorts (**Figure 2C**). Fold-change analysis revealed that multiple variable region peptides were significantly upregulated in individuals with PASC compared to convalescent controls (CONV). As shown in the volcano plot (**Figure 2C**), numerous IGHV, IGKV, and IGLV chains exhibited elevated expression, highlighting a broader shift in variable region usage or expansion of specific B cell clones in the PASC group. This differential abundance suggests persistent or skewed antibody responses that may contribute to the immunological landscape of PASC.

Overall, dysregulated plasma immunoglobulin patterns in PASC may reflect an ongoing B cell activation or an altered humoral immune response at the mucosal sites in individuals with persistent post-viral symptoms.

### PASC participants exhibit elevated levels of cTFH and MAIT cells in the blood

To further define dysregulation of the innate and adaptive immunological system in PASC participants, we performed high-dimensional profiling of PBMCs using CyTOF with a comprehensive 26-antibody panel targeting key cell lineage markers. To visualize cellular subsets, we performed UMAP clustering of concatenated samples from all individuals across both groups and all timepoints (**Figure 3A**). Within the CD45+ lymphocyte population, we identified clusters corresponding to canonical immune cell lineages, including total CD19+ B cells, CD3+CD8+ T cells and CD3+CD4+ T helper cells (including CD25hi CD127- T regulatory cells, Treg and CXCR5+ circulating T follicular helper cells, cTFH), unconventional T cells (mucosa associated invariant cells, MAIT and γδ-T cells), NK cells, innate-like lymphocytes type 2 (ILC-2), and myeloid cell subsets such as monocytes, myeloid dendritic cells (mDC), basophils, as well as plasmacytoid DCs (pDCs; **Figure 3A–B**). A similar clustering pattern was observed using t-SNE (**Figure S2**). As shown in **Figure 3B**, each cluster displayed characteristic marker expression consistent with lineage identities as defined in our gating strategy (**Figure S3**). When comparing the CD45+ immune cell subpopulations between the CONV and PASC groups at T0, T1, and T2 (**Figure 3C**), we observed differences in cluster size and density for cTFH and MAIT cells between groups, though these cell groups did not exhibit marked temporal changes. This prompted a more detailed subtype-specific analysis based on traditional gating (**Figure S3**).

**Figure 3.**
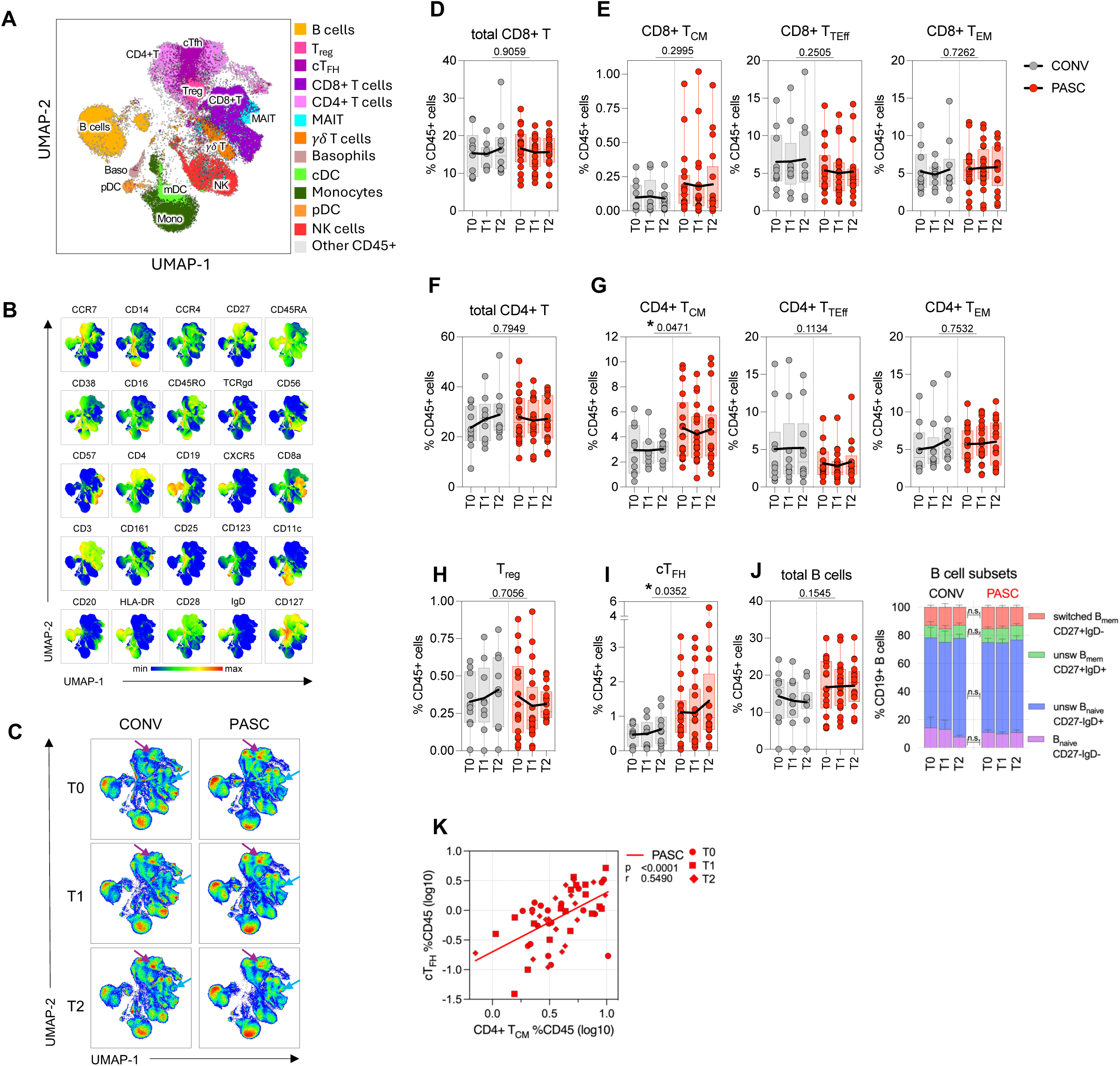
CyTOF analysis of PBMC reveals enrichment in CD4+ T_CM_ and cT_FH_ in PASC individuals. (**A**) Uniform manifold approximation and projection (UMAP) plot of clusters detected by CyTOF analysis of all CD45+ cell subsets within concatenated samples from all individuals in PASC and CONV cohorts at all timepoints. Distinct, color-coded cell subsets within the PBMC are overlayed on the total CD45+ population (grey) on the UMAP map. (**B**) Multigraph color maps of all markers used in cell phenotyping analyses; colors scaled from blue to red show min to max expression intensity. (**C**) Pseudo-color density UMAP maps of CD45+ cells from concatenated samples from CONV (left) or PASC (right) individuals at T0, T1 and T2 collection timepoint. UMAP clustering and cell types in **B and C** correspond to (**A**), color coded arrows indicated clusters with cT_FH_ (purple) and MAIT (blue) cells. (**D, E**) Frequency of total CD8+ T cells within CD45+ cells (**D**) and specific cell subsets within the CD8+ T (**E**): T_CM_-central memory, T_TEff_-terminal effector, T_EM_-effector memory. (**G**) Graphs represent total frequency of CD8+ T_CM_ (left), T_TEff_ (middle) and T_EM_ (right) within CD45+ cells. (**F**) Frequency of total CD4+ T cells within CD45+ cells. (**G**) Graphs represent total frequency of CD4+ T_CM_ (left), T_TEff_ (middle) and T_EM_ (right) within CD45+ cells. (**H** and **I**) Graphs represent total frequency of CD4+ T regulatory cells-T_reg_ (**H**) or peripheral, circulating follicular helper cells-cT_FH_ (**I**) within CD45+ cells. (**J**) Frequency of total CD19+ B cells within CD45+ cells (left) and cell subsets within the CD19+ B (bargraph on the right) based on the expression of CD27 and IgD. (**K**) Pearson correlation analysis between frequency of cT_FH_ and CD4+ T_CM_ within total CD45+ cell population. R and p values are calculated for Log10 transformed cell frequency numbers at T0, T1 and T2 timepoints for PASC patients. Each symbol represents a single individual at a specific time-point. In **D-J** graphs show Tukey analysis with min-max, each dot represents individuals in PASC (red), CONV (grey) cohorts, bar-connecting black lines show the mean values at distinct timepoints. In **D-J** numbers on the top of each graph show p values between experimental groups calculated for all timepoints calculated by 2-way ANOVA. Black (*) indicate significant difference *p < 0.05. n.s.-not significant (p > 0.05).

The frequency of CD3+CD4+ and CD3+CD8+ T cell populations within the total CD45+ PBMC did not vary significantly between the PASC and CONV groups over time (Figure **3D and F**). Similarly, within the CD8+ T cells, the CCR7hiCD45RA-CD45RO+ central (TCM) memory, CCR7lo/-CD8+CD27+ effector (TEM) memory cells and CCR7lo/- CD8+CD27- terminally differentiated effector (TEff) were similarly distributed in both experimental groups (**Figure 3E and Figure S4**). However, in accordance with the clustering analysis in **Figure 3C**, PASC individuals showed a significant increase in the proportion of CD4+CCR7hiCD45RA-CD45RO+ central memory cells (TCM) within all CD45+ lymphocytes when compared to CONV (**Figure 3G** and **Figure S4**). We also observed a substantial expansion in the population of the CD4+CXCR5+ circulating follicular helper cells (cTFH) in PASC participants in contrast to CONV (p=0.0352; **Figure 3I**) possibly indicative of an ongoing germinal center (GC) reaction and B cell activation, even though the total numbers of circulating B-cells or B cell subsets were not different between PASC and CONV (**Figure 3J**). The fractions of CD4+ TCM correlated strongly with cTFH subsets in PASC (p<0.0001, r=0.55 for PASC; **Figure 3K**).

To better characterize possible persistent, systemic inflammatory activation in PASC we validated innate cell subsets in the blood including cells of myeloid lineage, antigen presenting cells (APC) and innate-like lymphocytes. The frequency of total Lineage-HLA-DR+CD11c+ mDC or Lineage-HLA-DR+CD11c-CD123+ pDC populations within CD45+ lymphocytes remained constant between the two experimental cohorts (**Figure 4A**). Similarly, we did not detect significant changes in the magnitude of total CD14+ monocytes, nor any of the monocyte subpopulations (**Figure 4B**) or basophils (**Figure 4C**). Further, we characterized subpopulations of innate lymphocytes and invariant T cells (**Figure 4D-G**), and did not observe any significant differences between PASC and CONV cohorts in the circulating numbers of CD56+CD57+ mature, cytotoxic cells, circulating ILC-2 and the γδ-T cell populations (**Figure 4E, F**). In contrast, there was a significant 1.73-fold increase (p=0.0461) in the total frequency of CD3+CD4- CD28+CD161hi MAIT cells in PASC at all timepoints of specimen collection (**Figure 4G**). Importantly, the extent of increase in circulating MAIT cells in PASC participants was correlated with the high titers of anti-Env IgG at the group level (r=0.3, p<0.026; **Figure 4H**).

**Figure 4.**
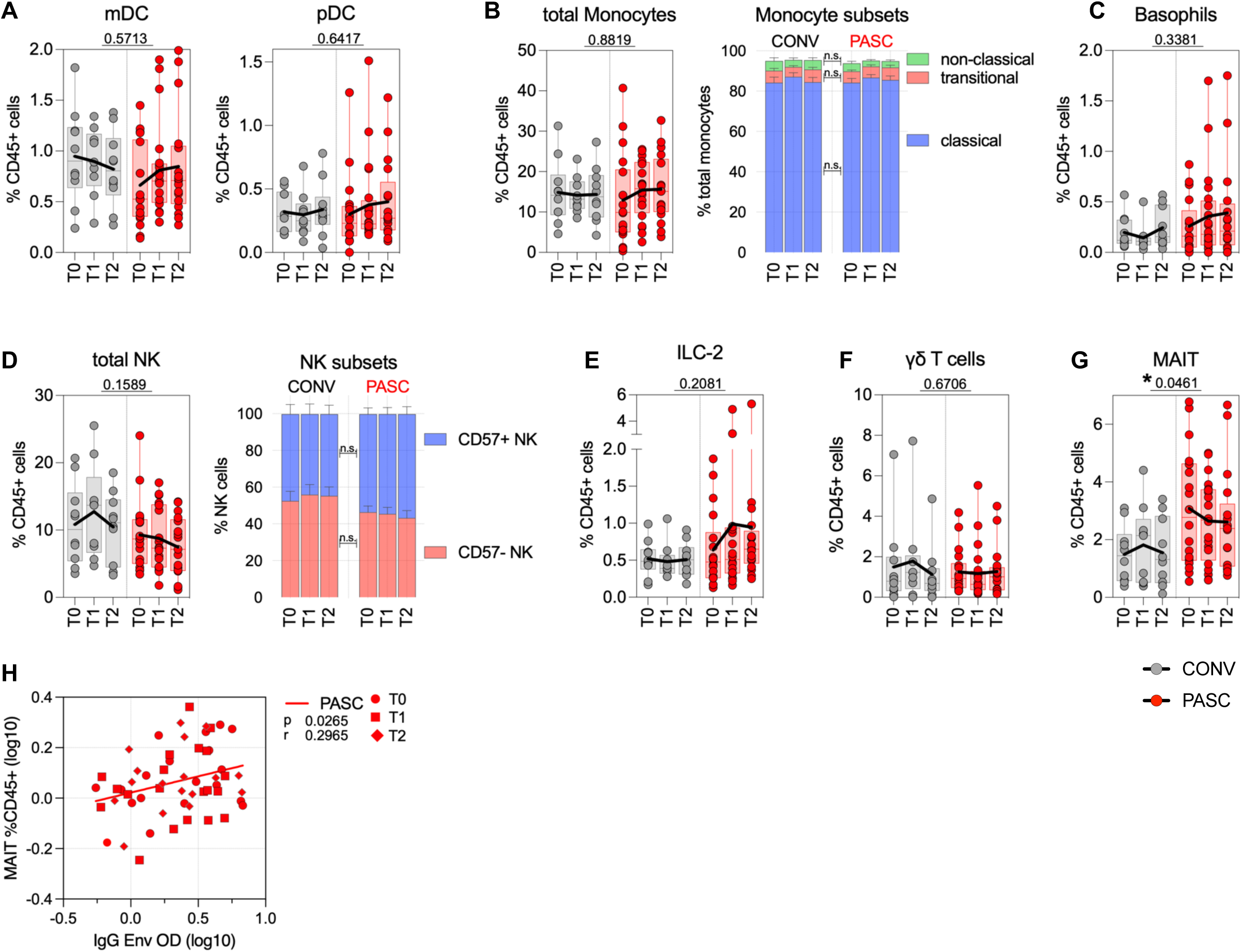
Elevated numbers of MAIT Cells in PASC Correlate with Anti-Envelope IgG Titers. **A-C:** analysis of myeloid innate cell subsets in CONV and PASC cohorts. (**A**) Graphs represent frequency of total DC subsets within CD45+ cells, CD11c+ mDC (left) and pDC (right). (**B**) Frequency of total monocytes (**B**) and Basophils (**C**) within CD45+ cells. In (**B**) bar graph on the right shows specific cell subsets within the CD14+ monocyte population: classical, transitional and non-classical monocytes. (**D**) Analysis of NK cell subsets. Total fraction of CD56+ NK cells within CD45+ cells (left), bar graph on the right shows specific NK cell subsets within parent population based on CD57 marker expression. (**E-G**) Graphs represent total frequency of ILC-2 cell (**E**), *γδ*-T cells (**F**) or peripheral, circulating MAIT cells (**G**) within CD45+ cells. (**H**) Pearson correlation analysis between frequency of MAIT cells and anti-Envelope IgG titers. R and p values are calculated for Log10-transformed cell frequency or IgG values at T0, T1 and T2 timepoints for PASC (red) group. Each symbol represents a single individual at a specific time-point. In **A**-**G** graphs show Tukey analysis with min-max, each dot represents individuals in PASC (red), CONV (grey) cohorts, bar-connecting black lines show means values at distinct timepoints. In **A-G** values on the top of each graph show p values between experimental groups calculated by 2-way ANOVA. Black (*) indicate significant difference *p < 0.05. n.s.-not significant (p > 0.05).

In summary, these analyses show an augmented number of circulating TFH cells that likely reflect an ongoing Germinal Center reaction and B cell activation. Additionally, the increase in MAIT cells in PASC and the positive correlation between circulating MAIT cell numbers and anti-Env IgG titers suggest ongoing mucosal inflammation in PASC individuals that may reflect persistence of viral antigens in the mucosal compartments.

### Inflammatory cytokine profile is associated with PASC

We analyzed serum cytokine milieu in samples from PASC and CONV individuals across all timepoints using a 96-plex cytokine panel (**Figure 5**). As shown in **Figure 5A**, a t-SNE analysis of cytokine profiles identified two distinct clusters corresponding to PASC and CONV cohorts, although samples from the specific timepoints (indicated by distinct symbols) clustered closely within each individual among the two groups. This suggested marked between-group differences in cytokine expression patterns, with relatively stable differences over time.

**Figure 5.**
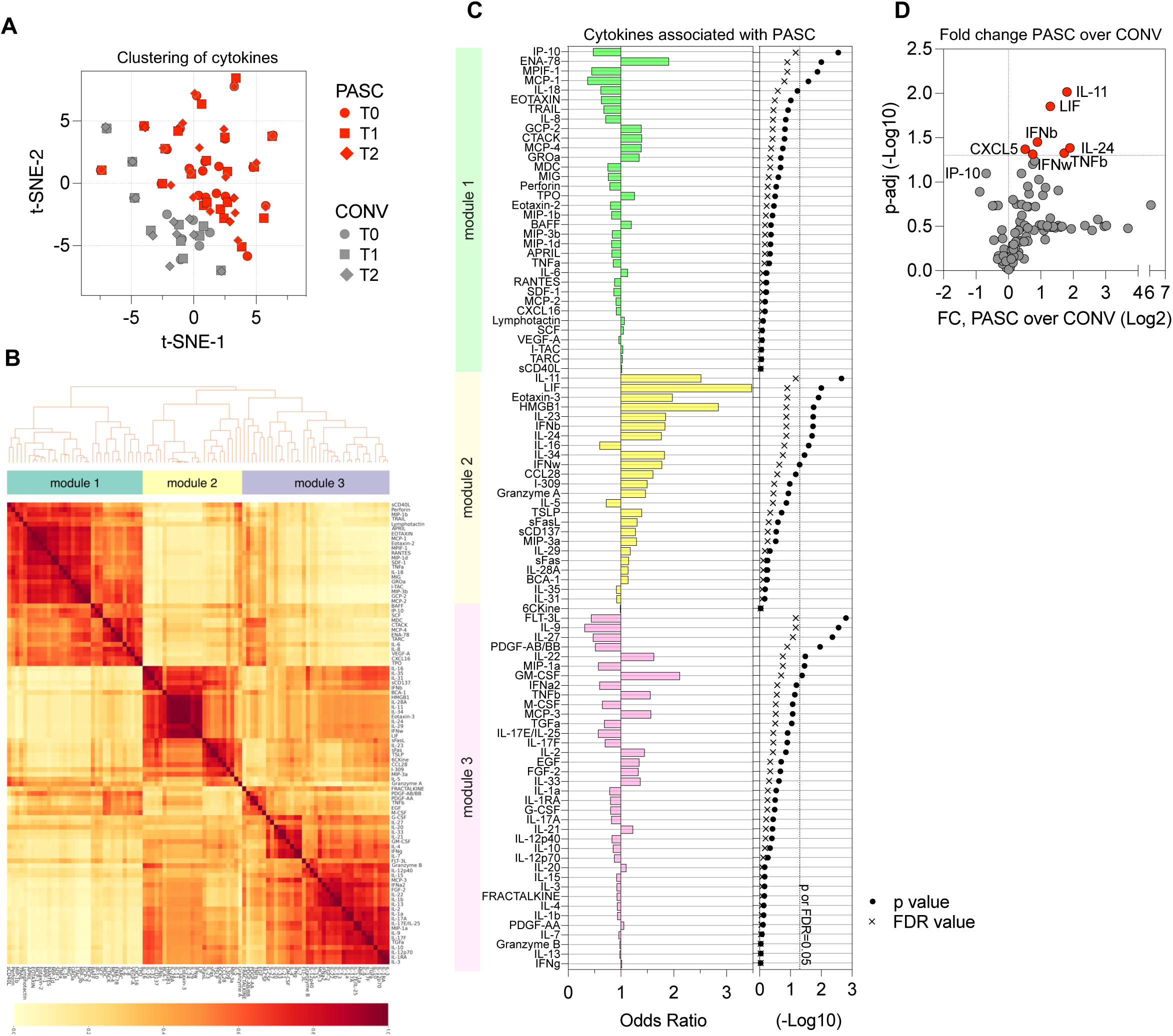
Distinct cytokine signatures are associated with PASC. (**A**) A t-distributed stochastic neighbor embedding (t-SNE) visualization of cytokine phenotypes between PASC (red) and CONV (grey), timepoints of plasma collections are represented as distinct symbols. (**B**) Heatmap of cytokine modules showing the cluster membership and reliability scores (i.e., the proportion of times that each pair of cytokines is allocated to the same cluster over 1000 random samplings of participant samples) of adjusted cytokines in all COVID-19 patients (both CONV and PASC cohorts) at all timepoints. (**C**) Logistic regression analysis of adjusted serum cytokine associations with clinical phenotype of COVID-19. Left panel represents Odds Ratio values for each cytokine or module as shown along the rows, grouped by their assigned module indicated in (**B**), bars show regression coefficient between cytokine or module level with PASC clinical phenotype. Right panel indicates p and FDR values assigned for each of the cytokines on the left. (**D**) Volcano plot shows fold-change of all 96 cytokines in PASC over CONV cohort. Dashed line at y-axis represent an adjusted p-value cutoff of 0.05. Significantly changed cytokines are marked in red.

To further explore co-expression patterns, we performed cytokine clustering analysis (**Figure 5B** and **Figure S5**). This approach revealed three major cytokine expression modules that were consistently observed across all individuals from both groups (**Figure 5B** and **Figure S5**). Each of the modules clustered a collection of cytokines with similar expression kinetics and values in both experimental groups over time. We calculated the odds ratios of cytokines in each of the modules to define the strongest association with PASC diagnosis (**Figure 5C**). A cluster of cytokines grouped in module 2 showed the highest odds ratio of positive association with PASC, with significant enrichment of cytokines such as IL-11, LIF, Eotaxin-3, IL-23, IFN-β, and HMGB-1, which were closely correlated with PASC outcome (**Figure 5C**). In module 1, expression of ENA-78 (CXCL5) was also positively associated with PASC whereas IP-10 (CXCL-10) correlated negatively with PASC diagnosis (**Figure 5C**). Finally, we compared changes in total cytokine values between PASC and convalescent cohorts over time (**Figure 5D**). Overall fold increase in circulating cytokine levels of PASC participants over the convalescent cohort over time was most significant for IL-11 and LIF, but also IFN-β, IL- 24, ENA-78 (CXCL5), and TNF-β, resembling cytokine modules defined by clustering analysis (**Figure 5B-D**).

The PASC diagnosis was strongly associated with a long-lasting systemic immune activation represented by overexpression of a several cytokines and chemokines which signal tissue damage and can potentially exacerbate chronic inflammation.

### Auto-IgG and -IgM antibodies in PASC highlight autoreactive response

Acute COVID-19 disease as well as the post-acute phase are often accompanied by an autoimmune reaction^19,40–42^. To assess for the presence of autoantibodies in PASC, we tested participant plasma for IgG and IgM reactivity against an array of over twenty-six thousand human proteins using HuProt technology. Global analysis of the protein array data showed an increase in total autoantigen reactivity in PASC participants when compared with convalescent (**Figures 6 and S6A**). We identified 16 proteins targeted by auto-IgG in plasma that showed significantly elevated titers in PASC compared to CONV serum, as illustrated in the heatmap of Z-scores representing the relative abundance of target-specific autoreactive IgG (**Figure 6A**) or top 20 proteins targeted by autoreactive IgM (**Figure S6A**). Mean absolute values of each of the auto-IgG titers remained stable with significantly higher relative intensity values in PASC individuals at least until the T2 timepoint (**Figure 6B**). **Figure 6C** visualizes the correlation matrix of the top 16 autoantigens targeted by plasma IgG autoantibodies in individuals with PASC identified in **Figures 6A and B**. Pearson analysis revealed that auto-IgG intensity values against 9 of the 16 autoantigens were strongly correlated (r > 0.7; p < 0.0001), suggesting that a subset of PASC individuals developed simultaneous autoreactive responses against multiple self-proteins. This pattern implied broad autoreactivity in a portion of the cohort, whereas other individuals exhibited more limited auto-IgG responses, targeting only one or a few specific autoantigens. The heterogeneity in autoantibody profiles highlights the diverse immunological manifestations of PASC and suggests multiple underlying mechanisms of autoreactivity. Interestingly, no single autoantibody was pathognomonic for PASC, and the targets of the autoantibody comprised a broad range of cellular targets, likely derived from different tissues.

**Figure 6.**
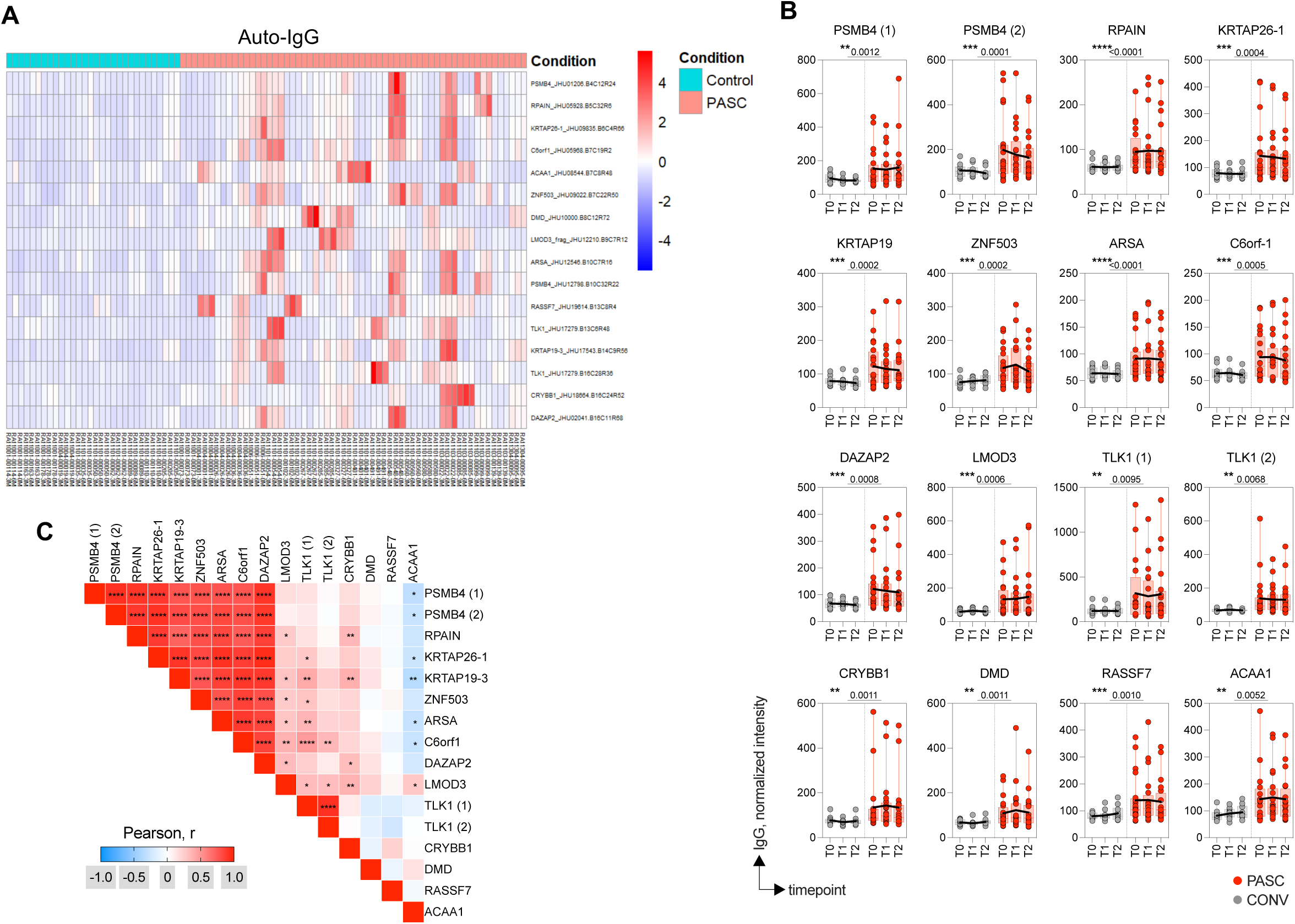
Autoreactive IgG repertoire in PASC highlights immune dysregulation. Autoreactive IgG titers were measured by HuProt protein array. **(A)** The heat map displays Z-scores representing the abundance of each top 16 target proteins of autoreactive IgG in PASC. Columns indicate individuals from PASC (red) and CONV (cyan) at each timepoint, rows represent HuProt array protein targets. (**B**) Absolute titers (normalized intensity) of auto-IgG against top 16 protein targets from (**A**) in PASC and CONV overtime. (**C**) Heat map with correlation matrix of autoreactive IgG titers (Log10) for distinct autoantigens. Pearson r-values are calculated for titers at T0, T1 and T2 timepoints for all PASC individuals. In **B** graphs show Tukey analysis with min-max, each dot represents individuals in PASC (red), CONV (grey) cohorts, bar-connecting black lines show means at distinct timepoints. Numbers on the top of each graph show p values between experimental groups calculated for all timepoints by 2-way ANOVA. Black (*) indicate significant difference *p < 0.05, **p < 0.01, ***p < 0.001, ****p < 0.0001.

Taken together, we detected a series of autoantibody titers upregulated in PASC participants and observed that they had developed IgG against multiple autoantigens.

### Effect of the Time Post-Infection on Antibody Responses in CONV and PASC

Finally, we evaluated whether the interval between the SARS-CoV-2 infection and the subsequent timepoints of specimen collection influenced the distinct dynamics of anti-SARS-CoV-2 antibody response defined by our analysis in both PASC and CONV participants. Participants were enrolled in the NIH RECOVER cohort over a period of 10 months between December 2021 and October 2022, as illustrated with the chronological timeline for each participant, indicating the date of infection, study enrollment, and each blood draw (**Figure 7A**). To standardize the timing gap across all individuals in both groups, we normalized all blood collection timepoints to the initial date of SARS-CoV-2 infection (**Figure 7B**). Despite the intra-cohort variability, there were no significant differences in the timing of specimen collection between PASC and CONV groups at any of the timepoints (T0, T1, and T2). To further investigate the dynamics of the antibody response, we stratified antigen-specific IgG titers by days post-infection for each individual in our experimental cohort. Spaghetti plot analysis exposed alterations in the temporal profiles of IgG responses to Spike, Envelope, Nucleocapsid, and Membrane proteins which varied overtime (**Figure 7C**). We performed Pearson correlation analysis between IgG titers against all SARS-CoV-2 antigens and the time interval from infection to specimen collection at each timepoint (T0, T1, and T2). This analysis revealed no significant correlation between the time gap and antibody titers at any of the measured timepoints, indicating that the interval between infection and blood draw did not significantly influence the magnitude of the antibody response (**Figure 7D**).

**Figure 7.**
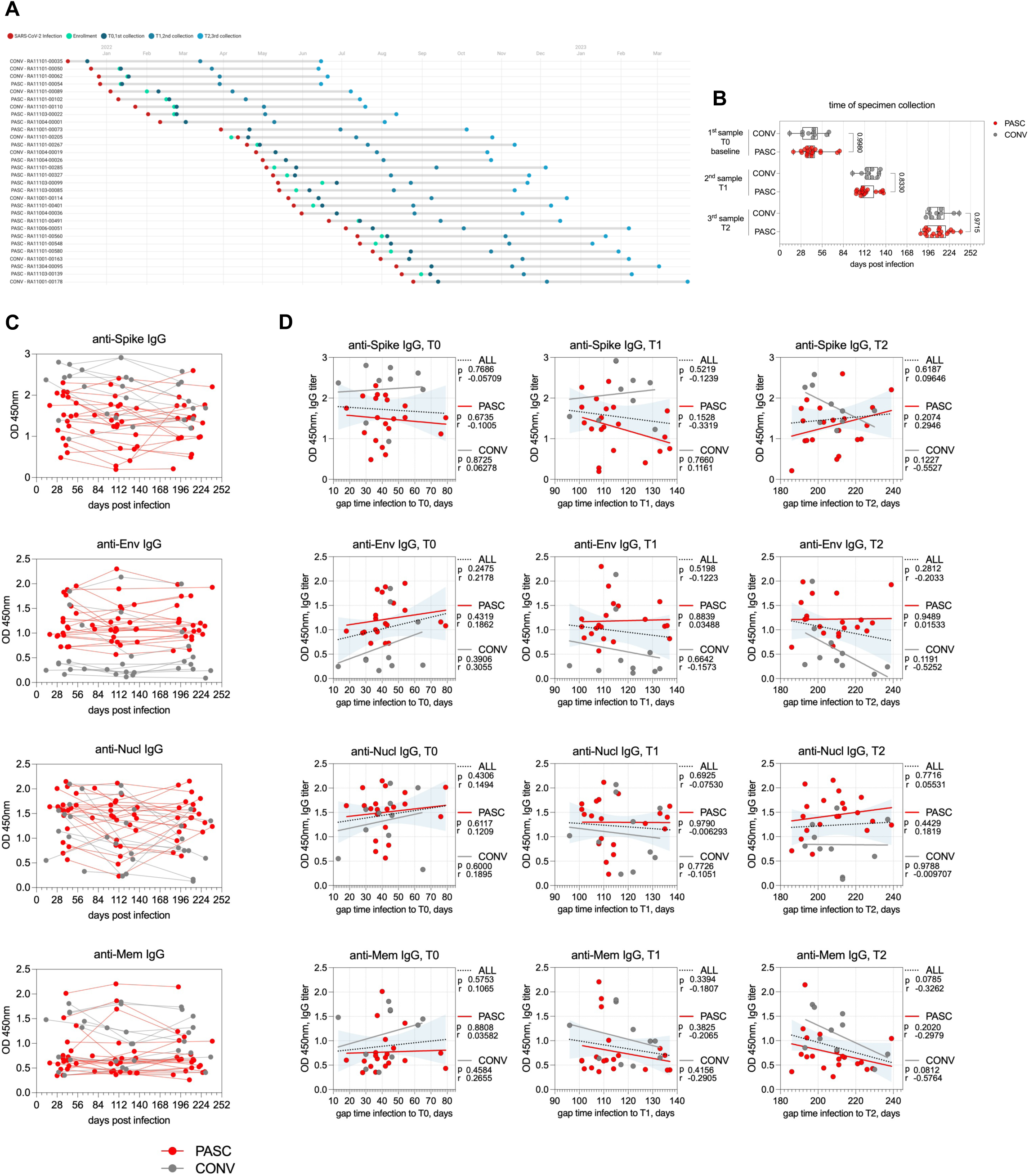
Effect of Time Post-Infection on Antibody Responses in CONV and PASC Groups. **(A**) A visual representation of a chronological sequence of infection, enrollment and blood specimen collection for individual participants enrolled in the study. Column on the left lists all CONV and PASC participants enrolled in the study, color-coded dots show timepoints fore infection, enrollment and each specimen collection. **(B)** Graph representing timepoints of each specimen collection at T0, T1 and T2 normalized to self-reported time of infection**. (C)** Spaghetti plots of trajectories of IgG titers for Spike, Envelope, Nucleocapsid, and Membrane measured on specific days post-infection. Each line represents individual participants, each dots show specific timepoint (days port infection) of blood collection and analysis. (**D**) Pearson correlation analysis between gap time from infection and each specimen collection (T0, T1 and T2) and IgG titers for Spike, Envelope, Nucleocapsid and Membrane. R and p values are calculated for IgG values at T0, T1 and T2 gap-to-infection timepoints for all participants (ALL, dotted line), CONV (grey) and PASC (red) cohorts. Each symbol represents a single individual at a specific time-point and colors represent the experimental group. In **B** graph show Tukey analysis with min-max, each dot represents individuals in PASC (red), CONV (grey) cohorts. In **B** values on the top of each graph show p values between experimental groups calculated by 2-way ANOVA.

## Discussion

Here, we studied the humoral, cellular and molecular indicators of a potential immune dysregulation over time in participants suffering from PASC and those who did not exhibit any persistent symptoms after a resolved SARS-CoV-2 infection. Our study shows a distinct shift in the SARS-CoV-2-specific antibody profile with dominant IgG responses to Envelope and Nucleocapsid viral proteins in individuals with PASC over the period of at least 5-7 months after the initial infection. Further analysis determined a close correlation of high Envelope-specific IgG titers with elevated proportions of circulating MAIT cells as well as an increase in the cTFH during PASC. Additionally, PASC participants exhibited an inflammatory cytokine signature and developed distinct autoreactive antibody responses.

Among the many hypotheses regarding the underlying mechanisms of PASC, a dysregulation of the immune system has emerged as a prominent pathogenic mechanism^2,6^. In our study the elevated anti-Envelope and anti-Nucleocapsid IgG response in PASC suggest an ongoing antigenic stimulation and suggests persistence of viral proteins or protein fragments in chronically ill individuals ^28,43^. Although all participants in both study cohorts reported a positive vaccination status (**Table 1**), the higher anti-Spike IgG titers observed in the CONV cohort may suggest a more efficient vaccine-induced response in this group^36,44^. This interpretation is further supported by our data of predominant Spike-specific IgG4 levels in CONV participants (**Figure 1D, E**), consistent with previous reports demonstrating high IgG4 class-switching of Spike-specific IgG response in individuals who received multiple doses of a mRNA vaccine^37–39^.

IgG4 inherently has an altered capacity to efficiently bind Fc receptors leading to reduced antibody-dependent cellular cytotoxicity (ADCC) and antibody-dependent cellular phagocytosis (ADCP) related events^45^. In contrast, IgG3 can effectively bind IgG-Fc-receptor and induce NK cell mediated ADCC^46^. Therefore, it is plausible that PASC patients experience more inflammation-prone events due to the persistence of elevated IgG3, while the control group having higher levels of the less inflammatory IgG4 is comparatively protected.

The overall IgG titer against Spike, albeit lower in the PASC group, remained stable over time for up to 7 months after acute infection in contrast to the convalescent group where Spike-specific IgG response diminished at later timepoints. Such a pattern may suggest a sustained immunogenic stimulus in PASC, potentially driven by persistent presence of viral antigens or delayed resolution of immune activation. The exact reason of high immunogenicity of Envelope and Nucleocapsid proteins (but not Spike or Membrane) in PASC remains unclear, but the consistency of IgG responses within our cohort supports the hypothesis of a prolonged Envelope and Nucleocapsid viral antigen exposure.

The precise location of the latent residual SARS-CoV-2 virus or viral RNA and the tissue serving as a reservoir for ongoing antigenic shedding cannot be ascertained from the analysis of circulating peripheral blood. Our mass spectrometry analysis of plasma proteome revealed elevated IgA and IgM immunoglobulin and J-chain peptides in PASC participants, along with elevated levels of Ig variable region peptides but none of the IgG subclasses (**Figure 2**). This supports the notion of *de novo* immune activation, especially at mucosal sites, and may suggest that intestinal and lung mucosa may serve as a location for viral persistence and sustained immune stimulation^18,47^.

Reactivation of other latent viruses such as Epstein–Barr virus (EBV) during or following SARS-CoV-2 infection has been increasingly recognized as a putative contributor to the immunopathogenesis of PASC^19,48,49^. Though our analysis focused on SARS-CoV-2- specific immune response, our proteomic analysis showing elevated plasma levels of IGHA, IGHM, and immunoglobulin variable region peptides may reflect mucosal antibody activity which is consistent with EBV’s tropism for the oropharyngeal and gastrointestinal epithelium^50^. Lytic EBV replication in epithelial cells induces the expression of pro-inflammatory mediators, including IL-6 family cytokines such as LIF and IL-11 (**Figure 5**), as well as BAFF, which may collectively promote local B cell activation and polyclonal expansion of mucosal antibody-secreting cells. These findings are consistent with the possibility that EBV or other herpesvirus reactivation, together with ongoing SARS-CoV-2 antigenic exposure, may contribute to the systemic immune alterations observed in PASC. ^48^.

Our immunophenotyping of PBMCs revealed significant alterations in individuals with PASC compared to convalescent controls, specifically in the frequencies of circulating TFH, TCM and MAIT cells. Elevated proportions of circulating TFH likely indicate their critical role in B cell activation, and the generation of high-affinity antibodies during the acute but also post-acute phases of the infection. The surge of cTFH and CD4+ TCM cells also suggests a persistent ongoing T-cell driven immune response and dysregulated B cell activity^16,51^. One possibility is that persistent antigen exposure, potentially due to viral protein reservoir or latent viral reactivation, sustains TFH cell differentiation and circulation. Alternatively, this could indicate altered trafficking or retention of TFH-like cells outside of secondary lymphoid tissues. Elevated frequencies of cTFH could contribute to prolonged or atypical antibody responses, including those observed in our proteomic analyses showing sustained E- and N-specific IgG titers in PASC, as well as elevated mucosal immunoglobulin signature in the plasma proteome.

Elevated presence of MAIT cells in peripheral blood during PASC is indicative of systemic immune activation, altered tissue trafficking, or an ongoing mucosal immune stimulation^52,53^. During acute SARS-CoV-2 infection, MAIT cells undergo significant perturbation, exhibiting activation during acute disease followed by numerical alterations and functional impairments^54–56^. Interestingly, in our study, PASC participants demonstrate sustained elevation of MAIT cell frequencies for at least 6-7 months post-acute infection. This persistence strongly correlates with heightened IgG titers against the SARS-CoV-2 Envelope protein, suggesting a prolonged immune activation state that is absent in convalescent individuals. In fact, in an animal model of SARS-CoV-2 infection, MAIT cells have been shown to provide help to B cells and enhance Ab response^57^. This raises important questions regarding MAIT cell involvement in the pathogenesis of PASC^58^. Their continued presence could indicate chronic inflammatory signaling, potentially mediated by IL-18 and IL-12 or other combination of inflammatory cytokines, leading to dysregulated immune responses. Moreover, the sustained anti-

Envelope IgG levels point to ongoing B cell activity, which may be influenced by MAIT cell-derived cytokines such as IFN-γ and IL-17. In chronic viral infections like HIV and hepatitis, MAIT cells frequently exhibit exhaustion phenotypes, contributing to disease persistence rather than resolution^52,59–61^. A similar mechanism may be at play in PASC, where sustained MAIT activation perpetuates inflammatory cascades, leading to lingering symptoms. The interplay between prolonged MAIT activity and persistent B cell activation necessitates further exploration, as it could provide valuable insights into the immune dysregulation characteristic of PASC. Investigating exhaustion markers and functional capacity of these cells in PASC patients may help clarify whether they are contributing to pathogenesis or attempting to resolve residual viral presence. Given their known roles in chronic viral infection, it is plausible that MAIT cells serve as mediators of long-term immune alterations in SARS-CoV-2 infection, warranting deeper mechanistic studies.

The presence and persistence of autoantibodies in a subset of PASC participants further underscores the immune dysregulation that we observed at the cellular level. Interestingly, the pattern of autoantibodies varied between the affected PASC participants, suggesting that there is no single autoantigen that promotes a state of persistent autoimmunity and immune dysregulation^19,41,62^. This heterogeneity in autoantibody responses may reflect the heterogeneity of endotypes in our PASC population as well as the fact that the tissue injury observed during the acute and post-acute phases of the SARS-CoV-2 infection can vary widely between individuals. The location of the tissue injury and the types of cells involved in the various manifestations of tissue injury may dictate which autoantibodies are generated, thus resulting in substantial variability between individual PASC participants.

One limitation of our study is the relatively small cohort sizes and the heterogeneity within the PASC group, both in terms of demographics and clinical presentation. PASC encompasses a wide spectrum of symptoms, and a larger-scale immunophenotyping study may be required to ascertain differences between distinct PASC endotypes. The divergency of PASC symptoms remains a central issue for many studies and addressing this complexity will require large, more stratified studies with sufficient power to explore associations between immune features and clinical phenotypes.

Our analysis focused on immune cell populations in peripheral blood as the primary source of study specimens. However, both our findings and other studies^16,18,24,25,27,54,63^ suggest that the most significant immune activation, dysregulation, and potential viral reservoirs likely reside in tissues, particularly mucosal sites. This limitation poses a challenge, as human tissue samples are difficult to access, making blood-based studies the most feasible approach despite their inability to fully capture tissue-specific immune dynamics. While peripheral blood provides valuable insights into systemic immune responses, it may not reflect the localized inflammation and persistence of viral antigens in tissues.

While our approach is a limitation for the purpose of mechanistic analyses and determination of possible tissue reservoirs of viral persistence, the detection of persistent antiviral IgG and of key markers of immune dysregulation in peripheral blood has significant value for its implementation in the clinical setting because blood is readily accessible during routine outpatient visits. Our work highlights the potential value of obtaining longitudinal samples over a period of several months to aid in the diagnosis and monitoring of PASC in clinical settings and intervention studies.

In summary, our longitudinal immunoprofiling reveals persistent humoral, cellular, and molecular immune dysregulation in individuals with PASC. We demonstrate that IgG antibodies in PASC participants exhibit persistent skewing in SARS-CoV-2 antigen recognition, a pattern that correlates with long-lasting activation of circulating T follicular helper (cTFH) cells and mucosal-associated invariant T (MAIT) cells. Importantly, these features persist for at least six months following PASC diagnosis, consistent with a model of ongoing viral or antigenic persistence, possibly in the mucosa. This immunological imprint is further characterized by continued systemic elevation of inflammatory cytokines and the presence of autoreactive antibody responses, suggesting sustained immune activation and dysregulation. Taken together, our findings provide new insights into the immunopathology of PASC and support the hypothesis that unresolved antigen exposure may underlie the chronic immune perturbations that define this syndrome. These findings highlight the importance of longitudinal profiling in PASC patients for diagnostic purposes, and they also suggest that such longitudinal profiling could be invaluable for the monitoring of the efficacy of novel therapeutics in PASC or Long COVID.

## Acknowledgements

This research was funded by the NIH OT2HL161847, as part of the Researching COVID to Enhance Recovery (RECOVER) Initiative, with subawards to the Illinois Research Network (subaward ADU-14-21; J.A.K., B.S.P., and J.R.) and NIH RECOVER Pathobiology Research Program (NIH OT2HL161847 subaward PATHO-PH2-SUB_22_23 to B.S.P. and J.R.). The content is solely the responsibility of the authors and does not necessarily represent the official views of the RECOVER Initiative, the NIH, or other funders. Authorship has been determined according to ICMJE recommendations.

## Disclosures

Jerry A. Krishnan, MD, PhD, reports grants from National Institutes of Health during the conduct of the study; research grants from the American Lung Association, BioVie Pharma, COPD Foundation, and Patient Centered Outcomes Research Institute outside the submitted work; and personal fees from AstraZeneca, Inogen, MedImmune, RespirAI, and Verona Pharma.

## Methods

### Study design and specimens

This retrospective longitudinal study included samples collected from n=30 adult participants already enrolled in the parent Researching COVID-19 to Enhance Recovery (RECOVER) Adult Cohort Study ^33,35^. Participants in the parent RECOVER Adult Cohort Study were enrolled at 83 sites across the United States recruited through clinician referrals, mailings to participants who tested positive for SARS-CoV-2, or responses to medical center paper postings, websites, and advertisements. Adult participants were enrolled in this cohort approximately 6 weeks after symptomatic, polymerase chain reaction (PCR)–confirmed SARS-CoV-2 infection between December 2021 and September 2021. The parent RECOVER-Adult protocol was approved by the IRB at NYU Grossman School of Medicine and collaborating sites, including the University of Illinois at Chicago (UIC, IRB study #2021-1287). Cryopreserved PBMC, plasma and serum specimens were received from the RECOVER cohort at Mayo Clinic, Rochester. Cryopreserved PBMC were stored in liquid nitrogen, and plasma and serum were stored at -80 °C until use. Trained personnel processed and tested frozen specimens at UIC under BSL-2 or BSL-3 conditions. Details about participant cohorts are summarized in Table 1. Specific dates for the participant’s infection (self-reported), study enrollment, and each of the 3 collections are summarized in **Figure 7A**; the timepoints of specimen collections are summarized in **Figures 1A** and **7B** as days post-infection.

### Antibody ELISA

We used 96-well flat bottom microtiter ELISA plates (MaxiSorp Thermo Scientific Nunc Cat. No. 442404) pre-coated with 0.1 µg/ml of SARS CoV-2 B.1.1.529/Omicron recombinant Spike trimer protein (His Tagged, Acro Biosystems, Cat.No. SPNC52Hz), SARS-CoV-2 B.1.1.529/Omicron Membrane Glycoprotein (RayBiotech Code: 230- 01124-100) at 0.2 μg/ml, SARS-CoV-2 B.1.1.529/Omicron Envelope protein (ABclonal Cat.No. RP01263LQ) at 0.1 μg/ml, SARS-CoV-2 B.1.1.529/Omicron Nucleocapsid Protein (Ray Biotech Cat.No, 230-30164-100) at 0.1 μg/ml. The coating buffer consisted of 15 mmol/L Na₂CO₃, 35 mmol/L NaHCO₃, 7.7 mmol/L NaN₃ at pH 9.6, and the plates were incubated overnight at 4°C. The wells were washed three times with 300 µL/well of washing buffer (0.05% Tween-20 in PBS, pH 7.4) and dried before blocking. For blocking, each well was treated with 300 μl Blocking Buffer (2% BSA in Washing Buffer, pH 7.4) at room temperature for 2 hours. After blocking, 100 μl of heat-inactivated serum were added to the respective wells at dilutions: 1:3200 for Spike, 1:200 for Envelope, 1:2000 for Nucleocapsid and 1:200 for Membrane. The samples were diluted in sample dilution buffer (0.5% BSA in Washing Buffer, pH 7.4) and incubated at 37 ℃ for 1 hour. Following the serum sample incubation, 100 μl of Goat anti-human IgG polyclonal secondary antibody conjugated with HRP (Invitrogen Cat. No. A18805) Goat anti-Human IgA Polyclonal Secondary Antibody, HRP from (Invitrogen Catalog # PA1- 74395) or Goat anti-human IgG1, IgG2, IgG3 or IgG4 (all Invitrogen) was added to the respective wells and incubated at 37 ℃ for 1 hour. The antibody was diluted to 1:5000 in Antibody dilution Buffer. Next, 100 μl of the Substrate Solution was added to each well and incubated at 37 ℃ for 10 min. The Substrate Solution consists of 8 μl 3% H₂O₂ and 100 μl 10 mg/mL TMB in 10 mL of Substrate Solution A (50 mmol/L Na₂ HPO₄·12H₂O, 25 mmol/L Citric acid, pH5.5). Finally, 50 μl of stop solution one mol/L sulfuric acid was added to each well, and the plate was read in a microtiter plate reader at OD 450 nm to obtain the OD values for calculations.

### Milliplex cytokine detection

To track and correlate the progression of 96 different analytes, serum samples from 30 participants were collected across three time points, heat inactivated at 56°C for 60 minutes in a water bath, and frozen for storage. The samples were then thawed for use in a multiplex assay using a Magpix platform (Millipore Sigma) to quantify the concentrations of 96 different analytes using 2 different kits. Samples were processed and loaded onto multiple 96-well plates, with procedures in doing so adhering to the protocol prescribed in the “MILLIPLEX® Human Cytokine/Chemokine/Growth Factor Panel A Magnetic Bead Panel - 96-Well Plate Assay” and the “MILLIPLEX® Human Cytokine/Chemokine/Growth Factor Panel B Magnetic Bead Panel - 96-Well Plate Assay” guides packaged with the kits used in the assay. Kits used in the assay were provided by MilliporeSigma (HCYTA-60K-PXBK48 and HCYTB-60K-PXBK48). The reagents used in this assay, including standards, quality controls, and detection antibodies were included in the respective kits. To measure the concentrations of each analyte in the samples after well plate preparation, samples were run on the Luminex® MAGPIX® instrument, an instrument used for multiplexing assays, using the xPONENT® software. Calibration and performance verification were conducted, adhering to instructions in the kit guides, prior to running the samples. After running the samples, data was outputted onto a Microsoft® Excel® spreadsheet, which included average concentration and median fluorescence intensity measurements in each well.

### CyTOF

PBMC samples were retrieved from liquid nitrogen and thawed in a 37°C water bath for 1 minute or until the frozen media had dissolved. Subsequently, 1 ml of thawed PBMCs was transferred to 9 ml of cold RPMI-10% medium in conical tubes, and the mixture was allowed to equilibrate for 10 minutes. The PBMC-containing conical tubes were centrifuged at 1500 rpm at 4°C for 5 minutes. The supernatants were carefully removed, the cells were washed twice with RPMI-10% medium for 5 minutes each time, and the cells were suspended in 1 ml of RPMI-10% medium. PBMC viability was assessed using 0.4% Trypan blue. Following this, 2-3 x10^6 PBMC were mixed with 5 mL of Maxpar Cell Staining Buffer (CSB 201068) and centrifuged at 300 x g for 5 minutes. The cells were then treated with 5μL of Human TruStain FcX™ FcR (Biolegend Cat. No. 422301) blocking agent for 10 minutes, and 215μL of staining buffer was added at room temperature. Then, 270μL of the FcR-blocked PBMCs were placed into a 5 mL tube containing the dry antibody pellet containing metal-tagged antibodies from a 30-marker panel and incubated at room temperature for 30 minutes. Cells were washed with staining buffer, then fixed with 1ml of fixed solution (1.6% formaldehyde) for 10 minutes at room temperature. After another wash with staining buffer, cells were suspended in 1ml of intercalator Ir solution (Fluidigm Cat. No. 201192A) and incubated overnight at 4°C. The cells were washed twice with staining buffer and twice with 2ml of cell acquisition solution (CAS Plus 201248). Labeled PBMCs were loaded into the Fluidigm CyTOF2® instrument for data acquisition. FCS files were analyzed using FlowJo v10 software. Gating strategy is presented in Figure S3. For clustering analysis, 1x10^5^ CD45+ cells from each fcs file from each participant in the PASC and CONV cohort, at each collection timpoint were concatenated in a single fcs file for further analysis. We used FlowJo build-in analysis for UMAP and tSNE clustering.

### Virus neutralization assay

The serum from each sample was diluted at a 1:50 ratio to achieve a 3-fold dilution in DMEM Medium. 60 ul of SARS CoV-2 omicron virus (B.1.1.529) at 5.2x10^5 FFU/mL was mixed with the DMEM and then incubated at 37°C for 1 hour. The mixture was then added to the Vero E6 cells and incubated for another hour at 37°C. Subsequently, 2% Methylcellulose medium was added to each well, and the plates were incubated for 36 hours. The overlay medium was removed, and the plates were treated with 4% paraformaldehyde for 30 minutes, followed by 10% formaldehyde for 15 minutes. Virus-inactivated plates were taken out of the BSL3 facility and washed with PBS. Later, the plates were incubated with Polyclonal Anti-SARS coronavirus, guinea pig (BEI – NR0361) primary antibody mixed in perm wash at a concentration of 1:15,000 for 3 hours at room temperature. Subsequently, the plates were washed with PBS and incubated with Goat anti-guinea HRP-labeled antibody (Sigma – A-7289) at a 1:5000 dilution for 3 hours. After being rewashed, the plates were stained with 30ul of TrueBlue stain for 15 minutes at room temperature. Finally, the plates were read by an Immunospot plate reader, and 20-200 spots/foci were counted and calculated.

### Mass spectrometry

Sample Preparation:10μl of each were initially diluted with an equal amount of PBS and then mixed with 40μl of urea. From this dilution, 25μl of the sample was mixed with 5μl of urea and 200mM DTT. The mixture was then incubated at 56°C for 30 minutes. Following this, 3.3μl of 500 IAA was added to the sample and then incubated in a dark room for 40 minutes. Subsequently, 170μl of 100mM ABC and 8μl of Trypsin (2μg) were added to each sample, and the samples were incubated at 37°C overnight. The digestion process was stopped by adding 10% formic acid (FA) to reach a final concentration of 1% FA. The dried protein digests were resuspended in 400μl of 5% acetonitrile and 0.1% formic acid buffer for LC-MS analysis. The criteria for analysis included a 1% false discovery rate (FDR) for proteins and peptides, with a minimum of 1 peptide required for identification. A 1uL sample was analyzed using a Q Exactive HF mass spectrometer linked to an UltiMate 3000 RSLC nanosystem equipped with a Nanospray Flex Ion Source (Thermo Fisher Scientific). Digested peptides were loaded into a Waters nanoEase M/Z C18 trap column (100Å, 5um, 180um x 20mm) and then a 75 μm x 150mm Waters BEH C18 column (130A, 1.7um, 75μm x 15cm) and separated at a flow rate of 300nL/min. The LC solvent gradient was as follows: 5% B from 0-3 min, 8% B at 3.2min, 8-35% B at 110 min, 35-95% B at 119 min, washed 95% at 129 min, followed by 5% B equilibration until 140 min. Full MS scans were taken in the Q-Exactive mass spectrometer over the 350-1400 m/z range with a resolution of 60,000 (at 200 m/z) from 10 min to 130 min. The AGC target value for the full scan was 3.00E+06. The 15 most intense peaks with charge states 2, 3, 4, 5 were fragmented in the HCD collision cell with a normalized collision energy of 28%, and these peaks were then excluded for 30s within a mass window of 1.4 m/z. A tandem mass spectrum was acquired with a resolution of 30,000, and the AGC target value was 1.00E+05. The ion selection threshold was 1.00E+04 counts, and the maximum allowed ion injection time was 50 ms for both full and fragment ion scans. The spectra were searched against the UniProt human database using Mascot Daemon (2.6.0, updated on 09/28/21) with specific search parameters. The search results were then entered into Scaffold DDA software (v6.0.1, Proteome Software, Portland, OR) for compilation, normalization, comparison of spectral counts. The filtering criteria for protein identification were a 1% false discovery rate (FDR) for both proteins and peptides with a minimum peptide count of 1.

### HuProt assay for autoantibodies

HuProt™ arrays were used for the ImmuneProfiler Assay by CDI Labs. Both, arrays and serum samples were blocked for 1 hour prior to the assay. For blocking, serum samples were diluted 1:1000 into CDISampleBuffer while arrays were incubated with CDIArrayBuffer at room temperature with gentle shaking. Each blocked and diluted sample was then probed onto a HuProt™ microarray at room temperature for 1 hour with gentle shaking. After probing, the arrays were washed with TBST (1xTBS / 0.1% Tween 20) for 10 min X 3, and then probed with Alexa 647-anti-human IgG, Fc specific and Cy3-anti-human IgM at room temperature for 1 hour in a light-proof box with gentle shaking, followed by 3 washes with TBST for 10 min each and 3 rinses with ddH2O. The arrays were then dried with an air duster and scanned using a GenePix® 4000B scanner for data collection. Consistent reproducibility was noted among hits in the raw data obtained from the GenePix software within samples. Non-specific hits that directly bind to the secondary antibodies were eliminated from the analysis of the sample. CDI software was used to quantify the specificity of each individual sample to specific proteins on the array based on Z Scores. Further analysis on the HuProt data was performed in R. After background probes were removed the remaining 23,058 proteins were tested. Each time point was treated independently with the comparison of CONV (n=30) vs. PASC (n=60) between groups. T-tests for each protein were performed with the comparison between CONV vs. Pasc. To control for multiple hypothesis testing, we adjusted the p-value to reduce the false discovery rate using the Benjamini Hochberg method. We filtered significant proteins with the adjusted *p−value* < .05 and *(log fold change*) > .5.

### Analysis and Statistics

Statistical analysis of data was performed using GraphPadPrism v.10.2 software. Where indicated, datasets between PASC and CONV cohort were analyzed using 2-way ANOVA, p values between experimental groups were calculated for all timepoints. For Milliplex multiplexed cytokine profiles modular clustering analysis in Figure 5 we used R software with Cytomod^64^. We used Orange Datamining software v.3.3.8 for heatmap clustering analysis in Figure 2A. Data visualization in Figure 7A was done using Datawrapper. Figure 1A was created with help of BioRender.

---

**Figure S1.**
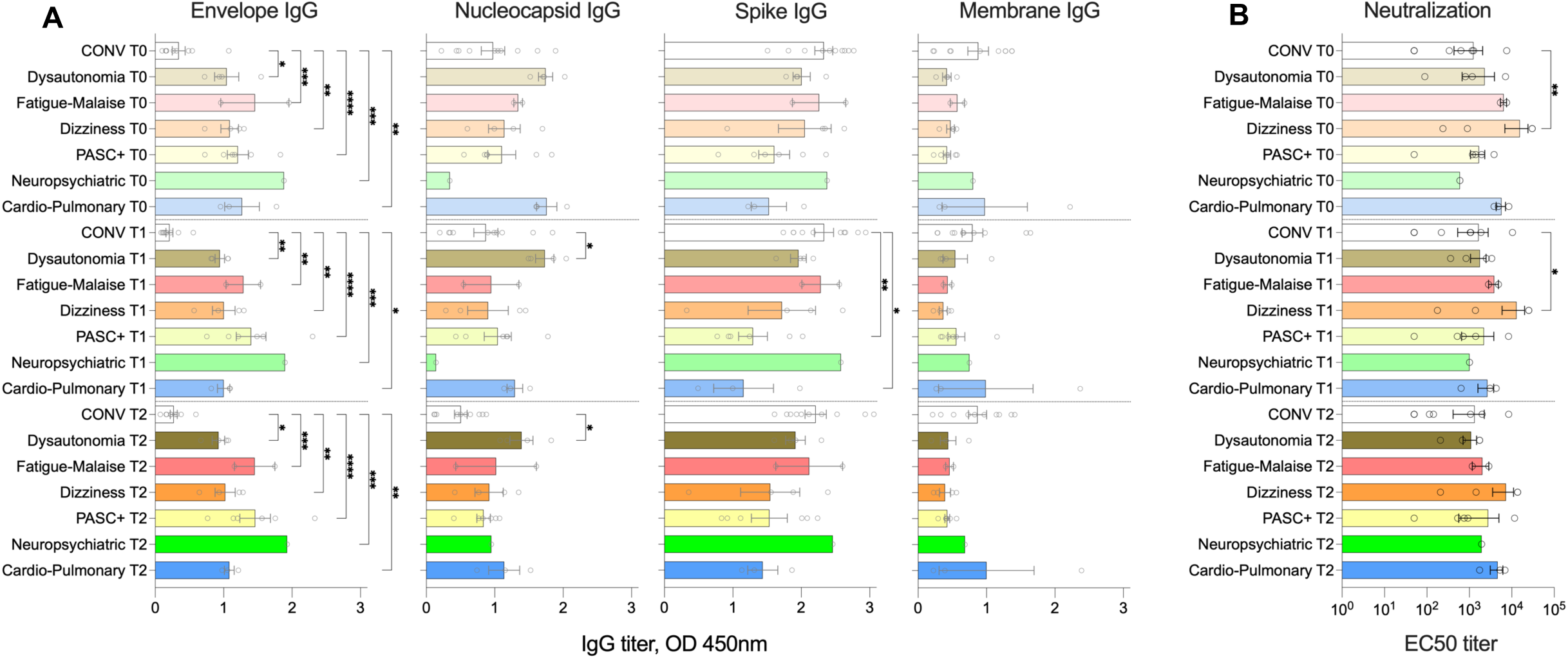
IgG titers in PASC patients sorted by symptom in comparison to CONV cohort. (**A**) Total IgG titers specific for Spike, Envelope, Membrane and Nucleocapsid as measured by ELISA at T0, T1 and T2 collection timepoint. X-axis represents ELISA OD values, bars show means +/-SEM, each dot represents participants in PASC, CONV (white bars). (**B**) SARS-CoV-2 neutralization titers are represented as EC50. p values between experimental groups calculated for all timepoints by 2-way ANOVA. Black (*) indicate significant difference *p < 0.05, **p < 0.01, ***p < 0.005, ****p < 0.0001.

**Figure S2.**
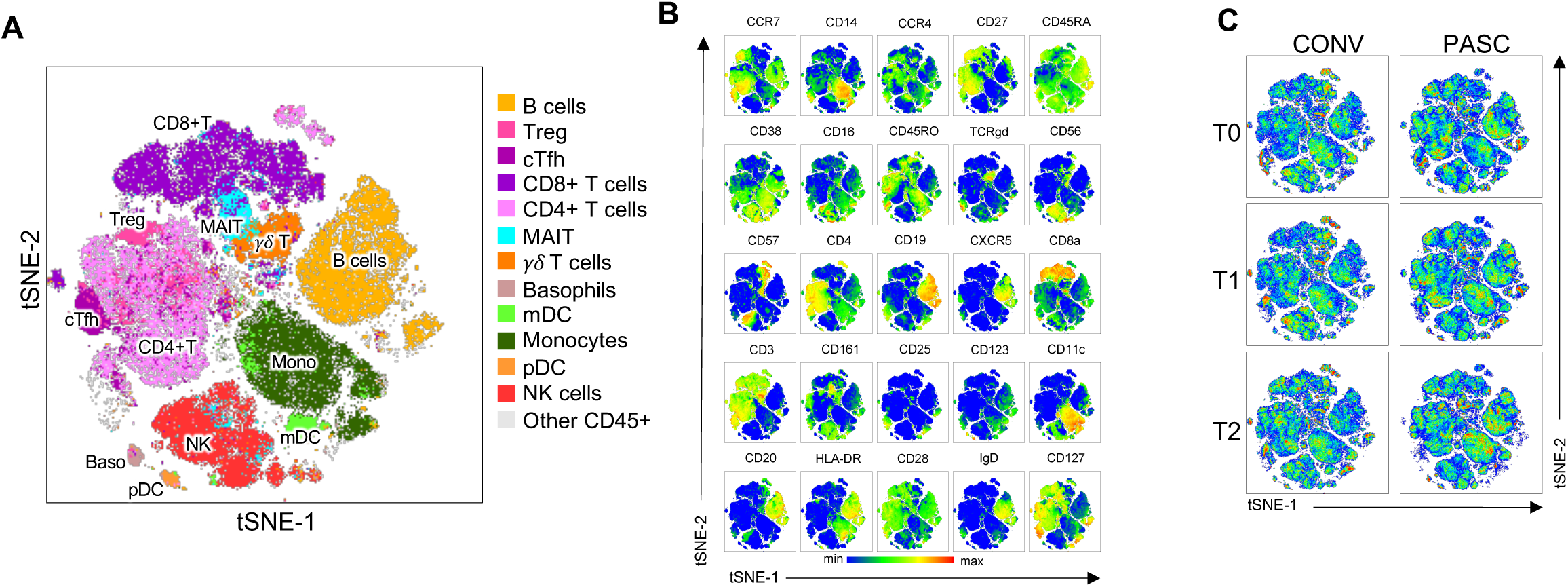
CyTOF analysis of PBMC by t-SNE. (**A**) A t-distributed stochastic neighbor embedding (t-SNE) analysis and map of all CD45+ cell subsets within concatenated samples from all participants in PASC and CONV cohorts at all timepoints. Distinct, color-coded cell subsets within the PBMC are overlayed on the total CD45+ population (grey) on the t-SNE map.(**B**) Multigraph color maps of all markers used in cell phenotyping analyses; colors scaled from blue to red show min to max expression intensity. (**C)** Pseudo-color density t-SNE maps of CD45+ cells from concatenated CONV (left) or PASC (right) cohorts at T0, T1 and T2 collection timepoint. tSNE clustering and cell types in **B-C** correspond to (**A**).

**Figure S3.**
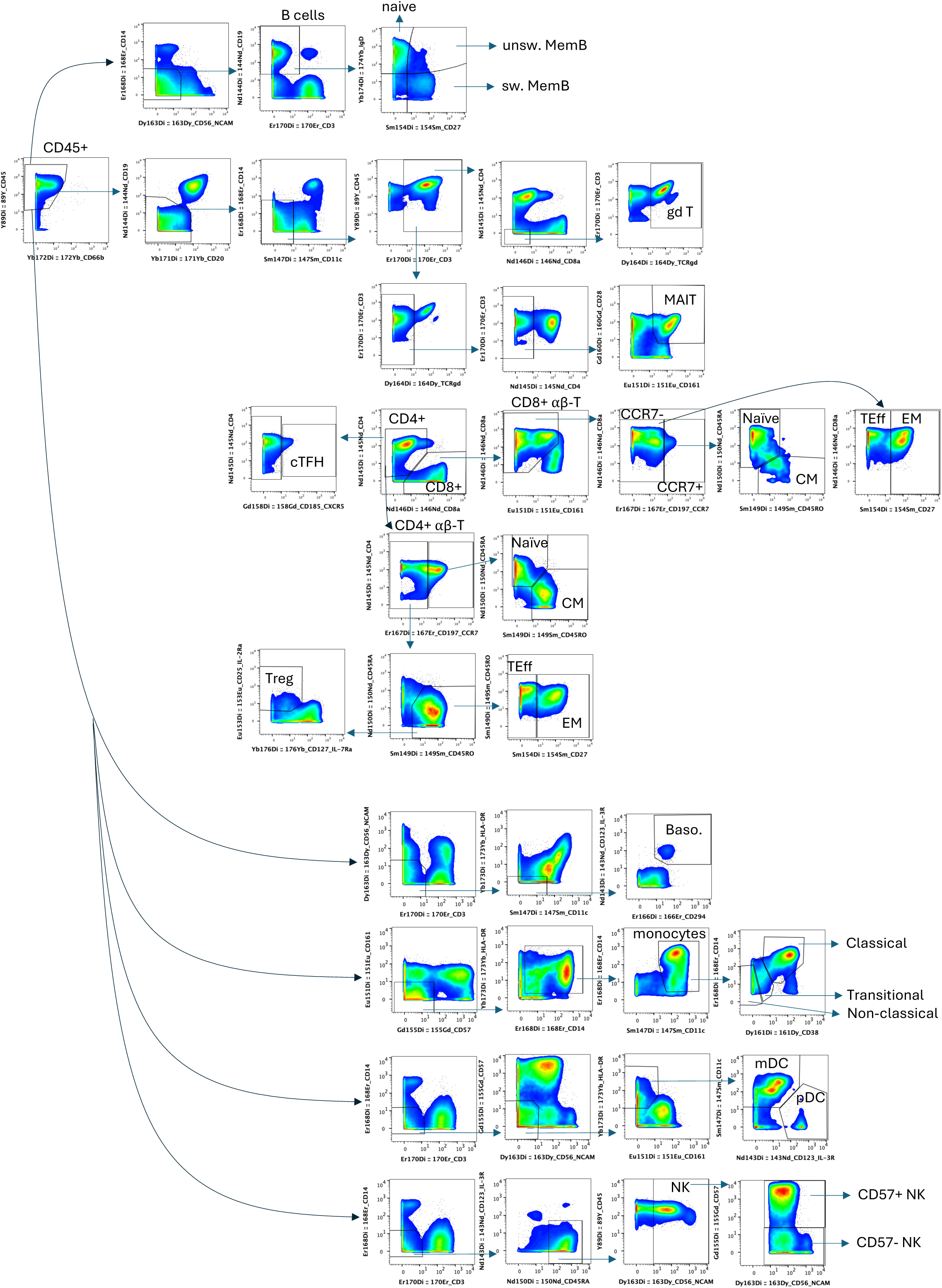
Gating strategy for PBMC populations.

**Figure S4.**
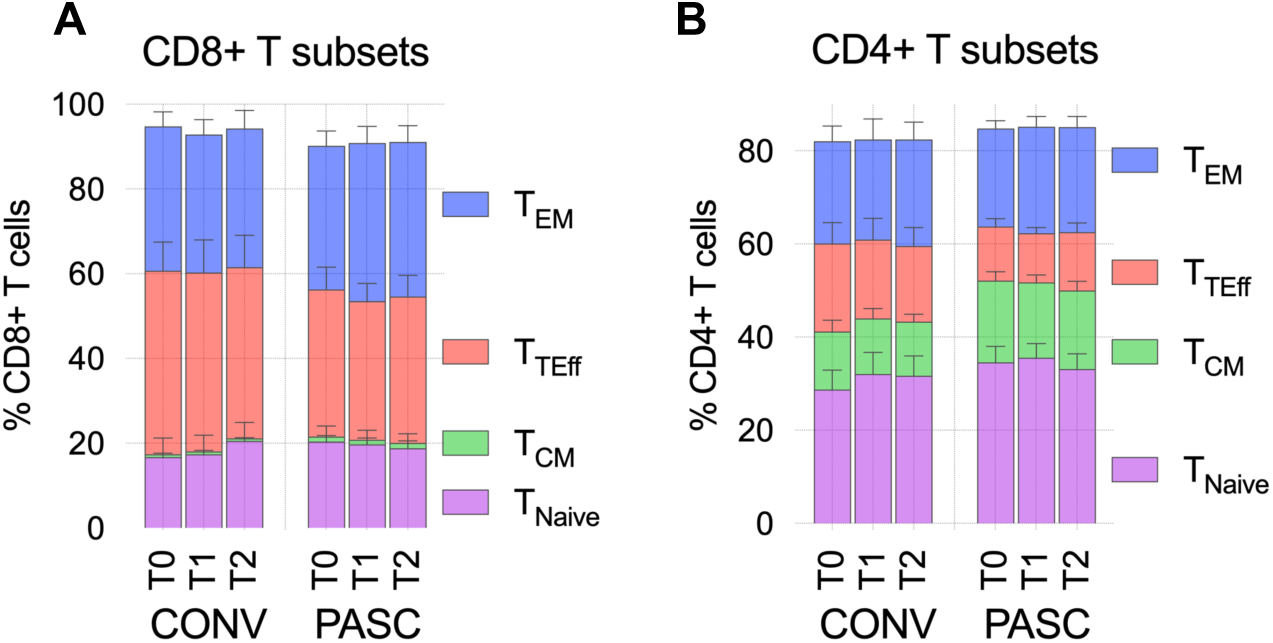
CyTOF analysis of T cell subsets. (**A and B**) Graphs represent frequency of CD8+ T_CM_, T_TEff_, T_EM_ and T_naive_ cell subsets within total CD8+ T cells (**A**) or CD4+ T cells. Cell subsets were defined as presented in the **Figure S3.**

**Figure S5.**
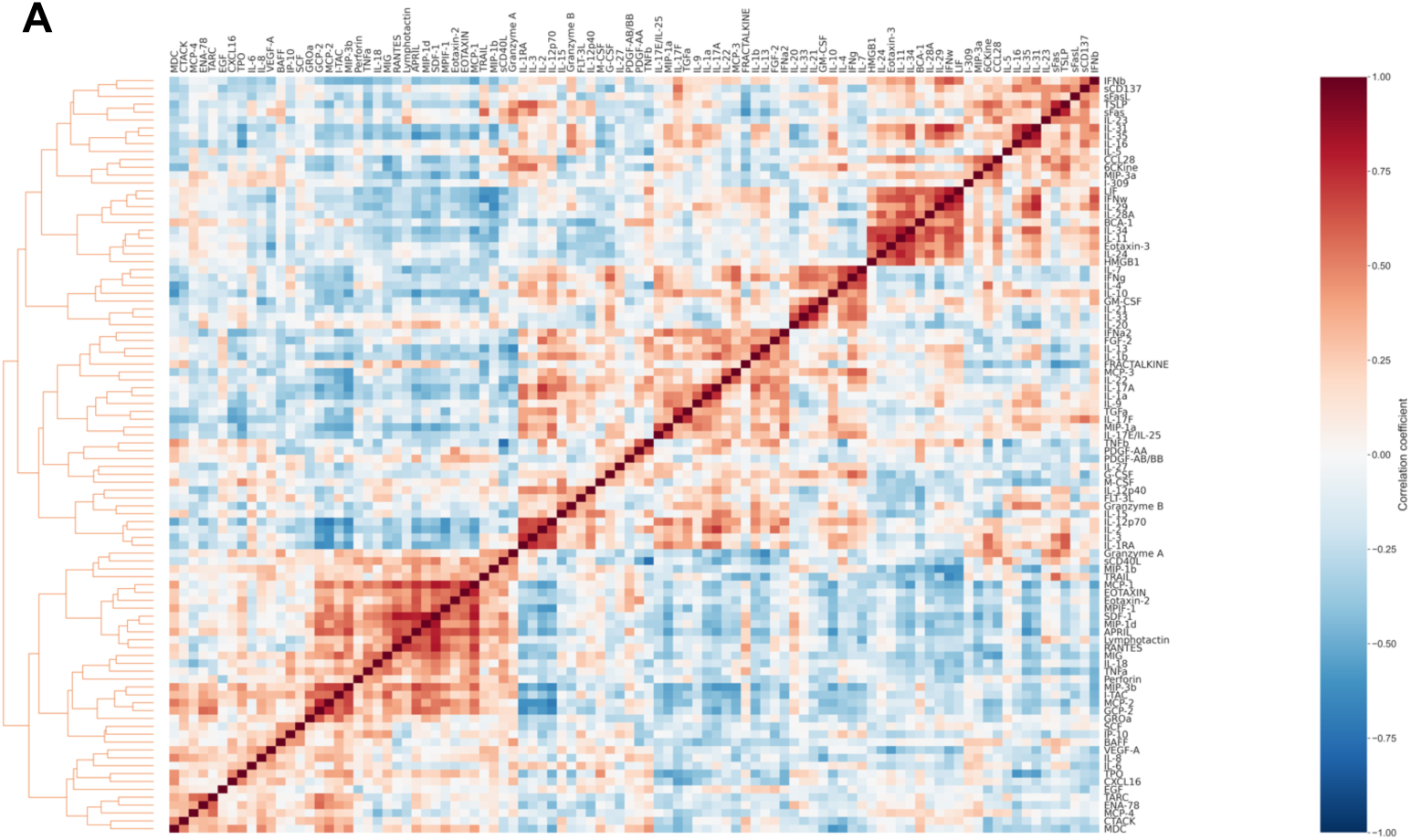
Cytomod analysis of cytokine signatures associated with PASC. (**A**) Pairwise Pearson’s correlations between cytokines following adjustment to the mean cytokine level.

**Figure S6.**
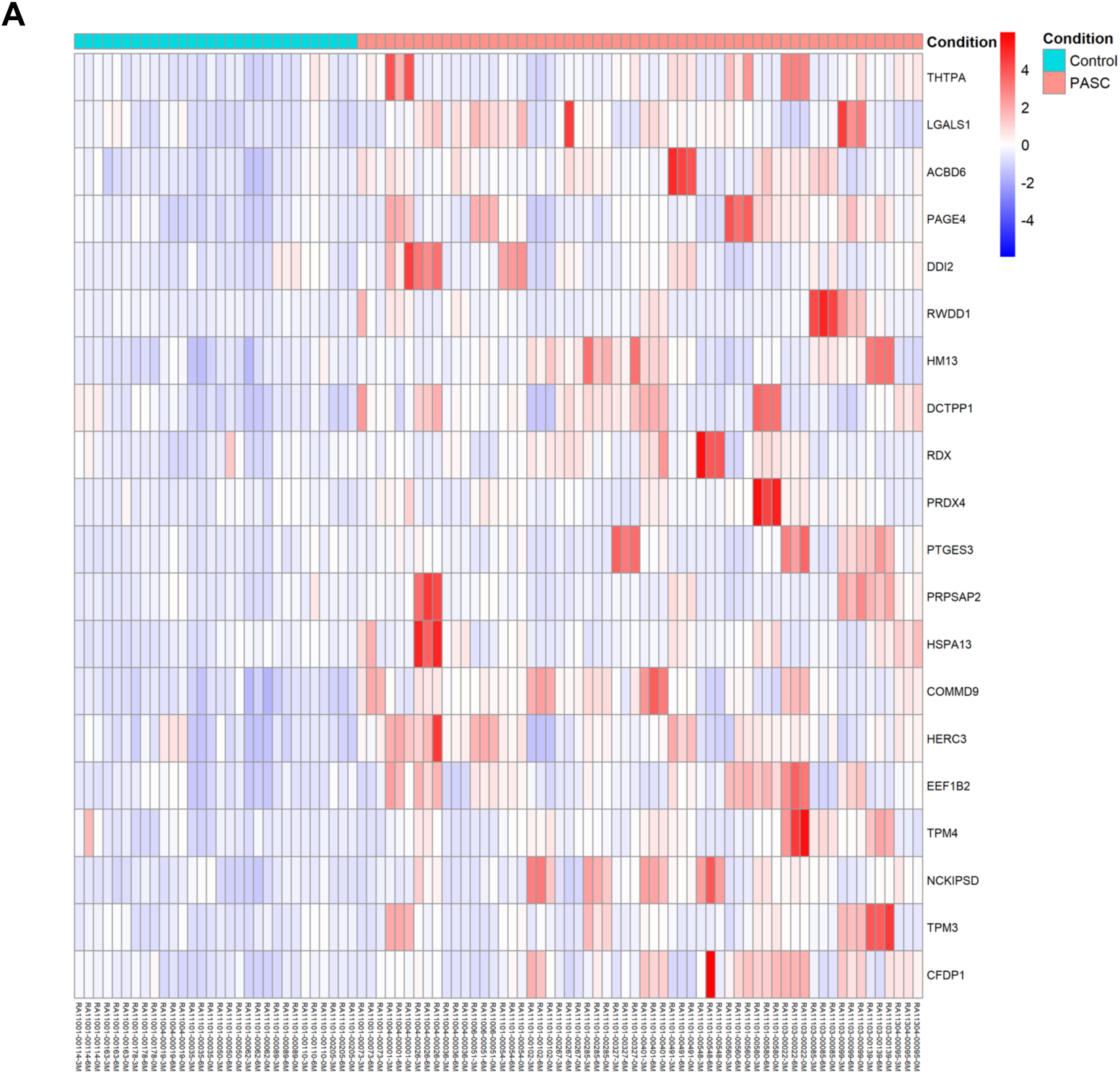
Autoreactive IgM repertoire in PASC. Autoreactive IgM titers were measured by HuProt protein array. **(A)** The heat map displays Z-scores representing the abundance of each top 20 target proteins of autoreactive IgM in PASC. Columns indicate individuals from PASC (red) and CONV (cyan) at each timepoint, rows represent HuProt array protein targets.

